# Alphaviral nonstructural protein-host RBP co-condensation as a mechanism to sustain virus replication

**DOI:** 10.1101/2025.11.30.691455

**Authors:** Yi Liu, Zhiying Yao, Qi Chen, Zemin Yang, Yexuan He, Xiaoxin Chen, Yun Zhang, Jiang Du, Jinjun Wu, Hong Joo Kim, Zhenshuo Zhu, Le Tian, Ziqiu Wang, Jing Huang, Yongdeng Zhang, Wenchun Fan, J. Paul Taylor, Peiguo Yang

## Abstract

It has long been recognized that the intracellular replication of alphaviruses critically relies on several key host RBPs, including G3BP1/2 and FXR1/FXR2/FMR1, but how these RBPs modulate alphaviral replication and whether it would be possible to target these RBPs for antiviral treatment are less explored. Here, using SFV as a model, we report that SFV nsP3 exploits G3BP for its condensation and transforms antiviral stress granules into proviral nsP3-G3BP co-condensates. The gel-like co-condensates of nsP3 and G3BP enrich and protect viral genomic RNAs from host RNase degradation and serve as viral translational hubs to promote viral replication. The mode of nsP3-RBP co-condensation is prevalent across alphaviruses, and disruption of nsP3 condensates is an efficient antiviral approach. Thus, these findings uncover a general anti-alphavirus strategy based on the conserved reliance of nsP3-RBP co-condensation.

## Introduction

Viral infection poses a significant burden on human health and remains a continuous threat to society. The frequent observation of viral inclusion body formation and large viral replication complexes suggests that the condensation of specific viral proteins inside host cells during the infection cycle is a prevalent phenomenon^1,2^. Liquid-liquid phase separation (LLPS) has been proposed as a mechanism for cellular compartmentalization^3,4^ and the formation of viral inclusion bodies, including those of Rotavirus, Ebolavirus, Measles virus, Influenza A virus, Vesicular Stomatitis Virus (VSV), Respiratory Syncytial Virus (RSV), SARS-CoV-2, and others^5–12^. Several recent studies have highlighted the importance of viral protein condensation in successful viral replication^13,14^. However, the host factors targeted by viral condensates and the interplay between virus and host interactions remain largely unexplored.

Semliki Forest Virus (SFV), a positive-stranded RNA virus of the alphavirus genus, serves as a model to investigate the significance of alphaviral structures during the infection cycle and to uncover specific host factors and pathways associated with alphaviral replication. Extensive evidence suggests that processes related to SFV are likely applicable to other viruses, particularly positive-strand RNA viruses^15^. Moreover, the alphavirus family includes numerous endemic members worldwide^15^, capable of causing severe diseases such as encephalitis and arthritis, with limited available antivirals or vaccines. Upon entering host cells, the positive-strand genome RNA functions as a template for the production of non-structural proteins (nsPs) using the host translation machinery^16^. The replication complex, comprising four viral nsPs, recruits host factors for optimal virus replication. The function of nsP3 is poorly understood but is shown to confer species specificity and influence host selection^17,18^. The C-terminal domain of nsP3 is highly divergent among alphaviruses and has evolved to interact with various host proteins^18–20^. It is unclear whether nsP3 could exploit specific host factors with this domain for viral replication and immune evasion, and if so, whether it would be possible to restrict viral replication via targeting such virus-host interactions.

In this study, we demonstrated that SFV nsP3 exhibits condensation ability and is essential for virus replication in cells. nsP3 condensates recruit G3BP1/2 through the C-terminal domain, thereby remodeling host SG dynamics and dampening antiviral SG formation. The nsP3-G3BP1 co-condensates protect genome RNA from decay and promote viral protein translation. Tandem G3BP1/2 binding motifs are prevalent in most Old World alphaviruses, while the New World alphaviruses have evolved to hijack other RNA-binding proteins (RBPs), such as FXR1/FXR2/ FMR1, for a similar purpose. We found that disrupting nsP3 viral condensate inhibits virus replication in human cells. This study reveals a conserved virus-host co-condensate formation in the alphavirus family, nucleated by viral nsP3, and suggests a potential targeting strategy for antiviral intervention.

## Results

### SFV nsP3 forms biomolecular condensate in human cells

SFV, a positive-strand RNA virus, encodes four non-structural proteins (nsPs), designated nsP1-4 (Figure S1A). Amino acid sequence analysis suggested the presence of intrinsically disordered regions (IDRs) in nsP1 and nsP3 proteins (Figure S1B). To investigate the expression pattern of each nsP in human cells, we constructed plasmids expressing each nsP fused with mCherry. Expression in human U2OS cells revealed that nsP1 and nsP3 form distinct foci in the cytoplasm, and nsP2 and nsP4 are diffusely distributed (Figure S1C). nsP1 contains a membrane-targeting signal, and the deletion of the transmembrane (TM) domain resulted in a diffuse pattern of nsP1 (Figure S1D), suggesting that nsP1 forms membrane-associated structure. Thus, we focused on investigating the nature of nsP3 structures in cells. Both gfp- and mCherry-tagged nsP3 could form foci in cells (Figure 1A). Fluorescence Recovery After Photobleaching (FRAP) experiment indicated a slow exchange dynamic of the gfp-nsP3 signal inside the foci, and live-cell imaging showed rare fusion between foci, suggesting that these foci exhibit gel-like properties (Figures 1B and S1E). The nsP3 foci remain stable upon treatment with 1,6-HD, indicating that hydrophobic interactions are not critical for their formation (Figure 1C). nsP3 foci in cells are distinct from membrane-bound structures, as revealed by the lipid-staining dye Dil (Figure 1D). A live recombinant SFV, with a gfp insertion in nsP3 proteins^21^, showed two types of nsP3 foci distinguished by Dil dye staining (Figure 1E). Large ring-like vesicles (Type I) are intermingled with membrane structures. A second type of nsP3 foci (Type II) tends to be smaller and distinct from membrane-bound structures (Figure 1E). Antibody staining of nsP3 in SFV-infected cells revealed the same pattern as gfp-fusion proteins (Figure S1F). FRAP of these smaller foci in live virus infected cells showed a similar slow dynamic (Figure 1F) and slow fusion (Figure S1G), as observed in cells expressing individual nsP3.

**Figure 1.**
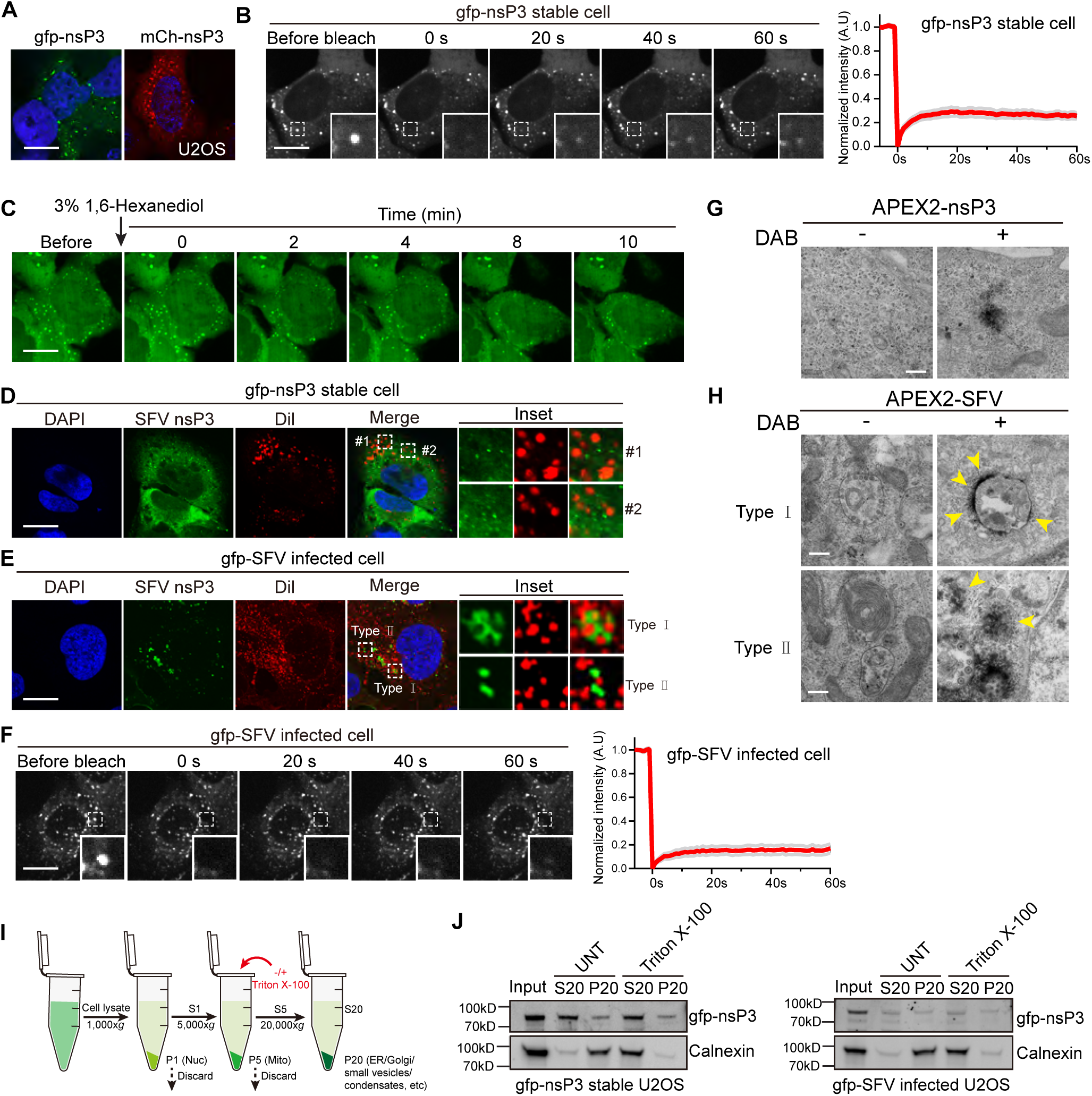
SFV nsP3 forms condensates in cells. (A) Representative image of gfp-nsP3 and mCherry-nsP3 expressed in U2OS cells. Scale bar, 10 μm. (B) FRAP experiment of gfp-nsP3 expressed in cells. Quantification curve on the right. Scale bar, 10 μm. (C) Representative images of 1,6-HD treatment of gfp-nsP3 expressing cells. Scale bar, 10 μm. (D) Representative images of co-localization between gfp-nsP3 and Dil labeled vesicles. Two insets are shown on the right. Scale bar, 10 μm. (E) Representative images of co-localization between gfp-SFV infected cells and Dil-labeled vesicles. Two patterns are detected: type I is membrane vesicle attached; type II is distinct from membrane structures. Scale bar, 10 μm. (F) FRAP experiments on gfp-nsP3 condensates in SFV-infected cells. Quantification curve on the right. Scale bar, 10 μm. (G and H) Representative EM images of APEX2-nsP3 stable expressing cells (G) and APEX2-SFV infected cells with or without DAB treatment (H). Two nsP3 patterns are observed: type I is membrane attached; type II is distinct from membrane structures. Scale bar, 500 nm. (I and J) Schematic diagram of biochemical fractionation assay (I) and WB results of the nsP3 fractions expressed in cells and during SFV infection (J). nsP3 was enriched in the pellet fraction of post-20,000 g centrifugation, and its pelleted distribution was unaffected by Triton X-100, a membrane-solubilizing detergent. In contrast, the membranous protein Calnexin was predominantly found in the supernatant upon treatment with Triton X-100.

To visualize the structure of nsP3 under electron microscopy (EM), we created an APEX2-nsP3 stable expression cell line and a recombinant APEX2-SFV virus. The virus successfully replicates in cells, as confirmed by biotinylation labeling and nsP3 staining (Figure S1H). We successfully identified non-membrane-bound structures in APEX2-nsP3 expressing cells (Figure 1G) and live virus infection (Figure 1H). Two populations of electron-dense nsP3 structures were identified in live virus infection: one associated with large cytopathic vacuoles (CPVs) and another for non-membrane-bound structures adjacent to CPVs in the cytosol (Figure 1H). We further performed a biochemical fractionation assay to determine if there is nsP3 in non-membrane-bound fractions. We found that a portion of nsP3 is insoluble both in the stable cell and during viral infection (Figures 1H and 1I). The proportion of nsP3 did not change with detergent treatment, suggesting that this fraction of nsP3 is not membrane-bound (Figures 1I and 1J).

Together, these results from fluorescent microscopy, APEX2-labeled EM, and biochemical fractionation assays demonstrate the presence of nsP3 condensates during live virus infection.

### nsP3 condensation is essential for SFV replication

To investigate the biochemical basis of nsP3 condensation, we created deletion mutants of nsP3 expressed in human cell lines (Figure 2A). nsP3 comprises an N-terminal Macro domain, a middle alphaviral unique domain (AUD) domain, and a C-terminal hyper-variable domain (HVD) (Figure 2A). AUD deletion completely abolished nsP3 condensation, and AUD alone was sufficient for aggregation in cells (Figure 2B), indicating an essential role of the AUD domain in mediating nsP3 condensation. We further mutated the two cysteines and three positively charged arginine residues inside the AUD domain of nsP3 (Figures 2A and S1I) and found that both the cysteine to alanine mutant (2CA) and the positive-charge abolishing mutants (3RA/3RE) completely abolished nsP3 condensation in cells (Figure 2B). The selection of charged residues was based on the observation that phase separation of the recombinant nsP3 *in vitro* is salt concentration-dependent (Figure S1J) and recent structural data of the AUD region from the Chikungunya virus in oligomerization^22^. SFV and Chikungunya virus belong to the same alphavirus family, and the AUD regions from both viruses showed high similarity in amino acid sequence (Figure S1I). The same effect in condensation disruption was also observed in the replicon constructs, which only contain the four nsPs from SFV (Figure 2C). We further performed IP experiment in WT 293T cells and found that AUD domain is required for mediating nsP3 self-interaction (Figure S1K). Considering that AUD alone can aggregate in cells (Figure 2B), these results collectively suggest the critical role of the AUD domain in mediating nsP3 condensation. To support the cellular findings, we purified recombinant nsP3 *in vitro*. We found that recombinant nsP3 protein can form condensates *in vitro*, whereas the AUD deletion, 3RE, and 2CA mutants all abolished the condensation of nsP3 *in vitro* (Figure 2D).

**Figure 2.**
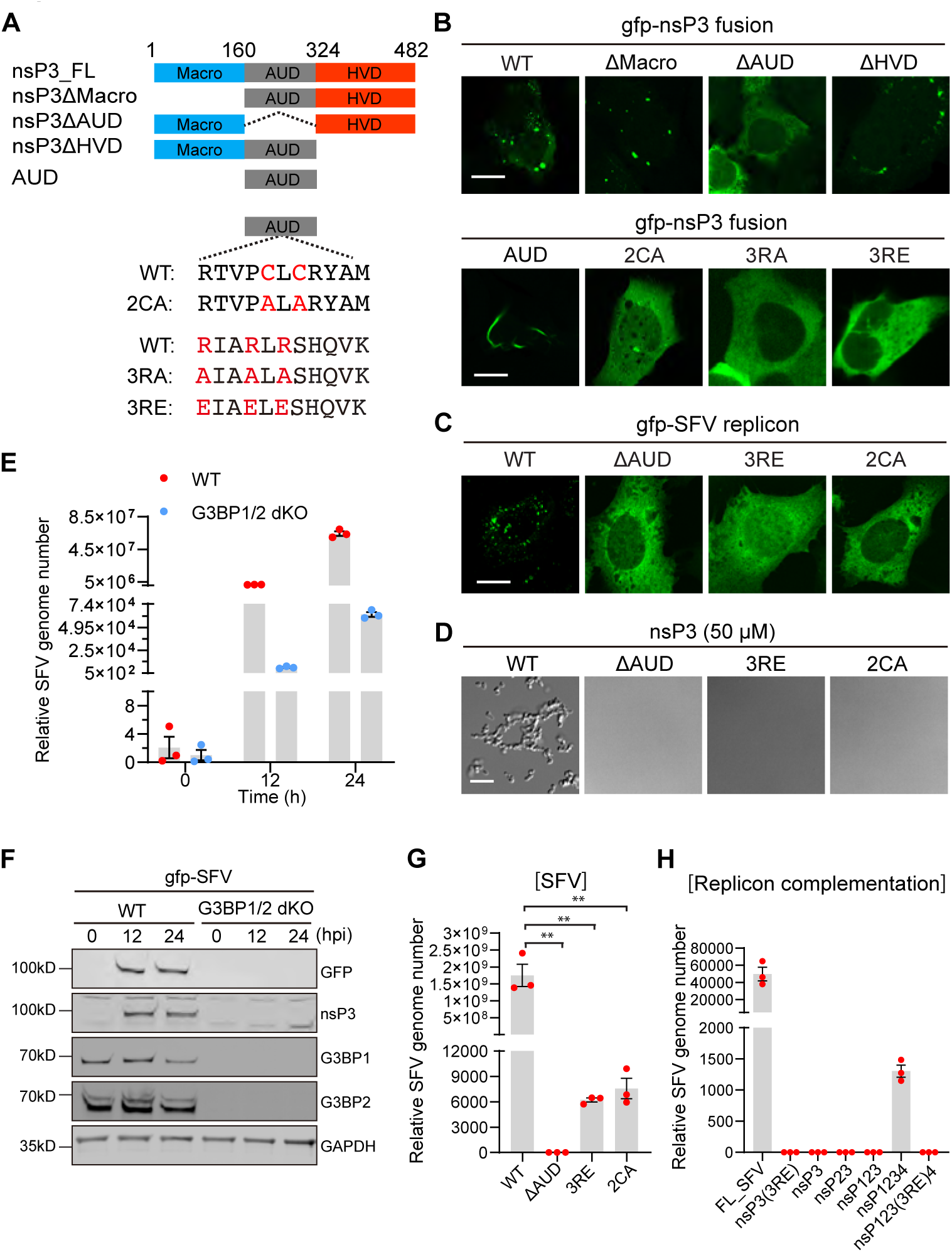
nsP3 condensation is required for efficient SFV replication. (A) Diagram of full-length SFV nsP3 protein and deletion or point mutations used in the study. (B) Representative images of WT and nsP3 mutants in cells. Scale bar, 10 μm. (C) Representative images of gfp-nsP3 replicon. The gfp is inserted in the HVD domain of nsP3. Scale bar, 10 μm. (D) Representative images of phase separation of recombinant nsP3 full-length and mutant protein *in vitro*. Scale bar, 20 μm. (E) Quantification of SFV replication by RT-qPCR in WT and G3BP1/2 dKO U2OS cells. Error bars indicate± SEM with n=3 replicates. (F) Representative WB images for nsP3 and G3BP protein expression during SFV replication. (G) Replication of SFV mutants in cells. Data are mean ± SEM; n=3 replicates. (H) Complementation of SFV(2CA) mutant by different replicons containing combinations of nsPs and mutations. SFV(2CA) mutant construct was co-transfected with each replicon construct, and the results were compared to those of FL_SFV. Data are mean ± SEM; n=3 replicates. Statistical significance was assessed in comparison to WT cells (E) or WT SFV (G) and (H) using an unpaired, two-tailed *t* test.

To study the role of nsP3 condensation in virus replication, we established an SFV replication assay in U2OS cells. We can recapitulate the strong reliance of SFV replication on G3BP1/2 proteins, observing a >1,000-fold reduction in viral replication in G3BP1/2 dKO cells compared to WT cells (Figure 2E). Consistently, no gfp-nsP3 protein expression was detected in G3BP1/2 dKO cells compared to WT U2OS cells (Figure 2F). To exclude the possibility of reduced viral entry in the dKO cells, we monitored the expression levels of SFV receptors, VLDLR and ApoER2/LRP8^23^. RT-qPCR results revealed no reduction in receptor expression in G3BP1/2 dKO cells (Figure S1L), suggesting that virus entry was not the primary cause of reduced virus replication in dKO cells.

To demonstrate the role of nsP3 condensation in viral replication, we created SFV mutants by introducing condensation-disrupting mutations, including AUD domain deletion, 3RE, and 2CA mutations. RT-qPCR results showed abortive infection of all three SFV constructs (Figure 2G). We further established a complementation assay by screening different nsP3-containing constructs. By co-transfecting SFV(2CA) and various nsP3-containing constructs, only a full-length replicon containing nsP1234, not other shorter fragments, could significantly rescue the 2CA mutant virus replication in cells (Figure 2H). The condensate-disrupting 3RE mutation abolished the rescue effect of nsP1234 replicon, underscoring the critical role of nsP3 condensation in mediating viral replication.

These results highlight the essential role of nsP3 condensation in SFV replication using both full-length virus constructs and a replicon complementation system.

### G3BP facilitates nsP3 condensation

Previous studies have shown the critical role of G3BP1/2 in alphavirus replication and spatial relationship with nsP3^17^. To dissect the role of SG-related host factors in nsP3 condensation, we tested other core SG network proteins, including TIA1, Caprin1, PABP, eIF4G, and eIF3η^24–26^. nsP3 cytoplasmic foci colocalize with G3BP1 and G3BP2 but not with other stress granule markers (Figure 3A). The same pattern was observed in recombinant SFV-infected cells (Figure 3B). Thus, the composition of nsP3 condensates was distinct from canonical SGs. Two-color super-resolution imaging further showed the intermingled distribution between G3BP1 and nsP3 inside the nsP3-G3BP co-condensate (Figure 3C), consistent with the ribbon-like structures previously observed in SG^27^. A similar intermingled pattern between nsP3 and G3BP1 was detected in SFV-infected cells (Figure 3D). These results implicate a specific G3BP-nsP3 interaction in nsP3 condensate formation in host cells.

**Figure 3.**
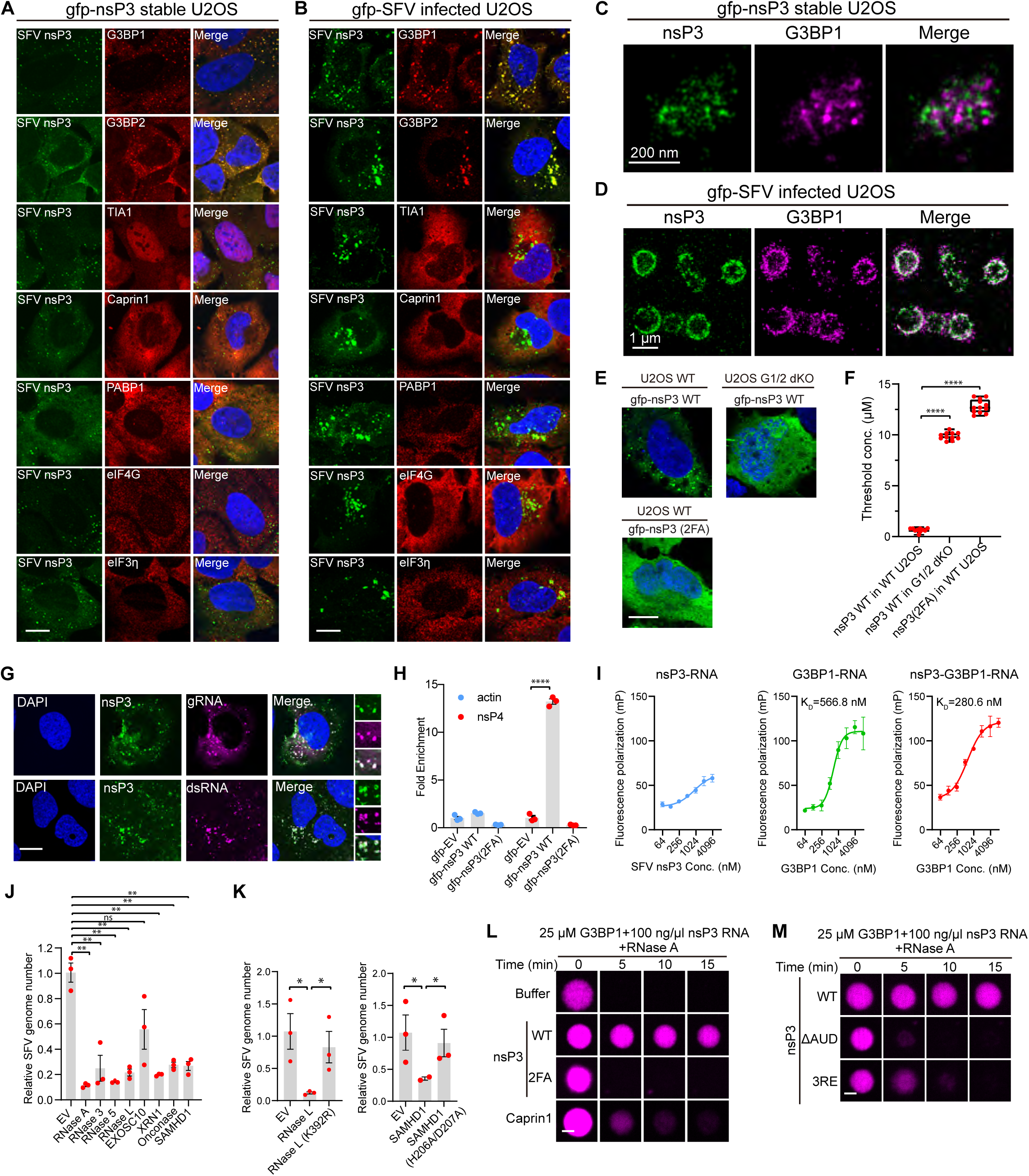
G3BP protein facilitates viral nsP3 condensation to shield viral RNA genome. (A) Representative images of localization patterns of SG markers with gfp-nsP3 in cells. Scale bar, 10 μm. (B) Representative images of localization patterns of SG markers with gfp-SFV infected cells. Scale bar, 10 μm. (C and D) Representative two-color super-resolution images of gfp-nsP3 with G3BP1 in individually expressed nsP3 cells (C) or SFV infection (D). Scale bars, 200 nm for (C) and 1 μm for (D). (E) Representative images of nsP3 mutant and nsP3 expression in WT and G3BP1/2 dKO U2OS cells. Scale bar, 10 μm. (F) Quantification of the threshold concentration for each condition in (E). Data are plotted as a minimum to maximum; lines indicate the first quartile (lower), median, and third quartile (upper) with n=10 values in the determined thresholding bin. See Methods for defining the bin. (G) Representative images of localization pattern between dsRNA and virus genome RNA with nsP3 condensates in SFV infected cells. An inset was shown on the right. Scale bar, 10 μm. (H) RNA-IP coupled with RT-qPCR to show the RNA binding feature of nsP3. Error bars are defined as ± SEM with n=3 replicates. (I) Fluorescence polarization assay to measure the RNA binding of G3BP1 and its enhancement by nsP3. Data are mean ± SEM with n=3 replicates. (J) RT-qPCR results show the viral inhibition effect of different RNase tested in cells. Data are mean ± SEM with n=3 replicates. (K) RT-qPCR analysis of the antiviral effects of the WT RNase L, SAMHD1, and their respective RNase-deficient mutants. Data are mean ± SEM with n=3 replicates. (L) RNA protection effect of nsP3 condensate using *in vitro* phase separation assay. *In vitro* transcribed SFV nsP3 RNA was labeled with Cy5-UTP, and the RNA signal was used to indicate the decay of RNA and dissolution of condensate after RNase A addition. Scale bar, 5 μm. (M) RNA protection assay of nsP3 mutants as in (L). Scale bar, 5 μm. Statistical significance in (F), (J), and (K) was assessed using an unpaired, two-tailed *t* test. Significance in (H) was assessed using a one-way analysis of variance (ANOVA) followed by Turkey test.

To test the role of G3BP proteins in nsP3 condensate formation, we transfected nsP3 construct into G3BP1/2 dKO cells and found that SFV nsP3 could barely form condensates in cells (Figure 3E). nsP3 interacts with G3BP1 via a tandem FGDF motif, and mutation of two critical phenylalanine residues in nsP3 abolished nsP3-G3BP interaction^28^. We further introduced nsP3(2FA) mutations to disrupt nsP3-G3BP1 interaction and found a similarly reduced ability to form condensate in WT cells (Figure 3E). We quantified the threshold concentration of nsP3 required for condensation in WT and G3BP1/2 dKO cells. The threshold concentration for WT nsP3 protein to form condensate is below 1 μM in WT U2OS, and this threshold increased about 10-fold in G3BP1/2 dKO cells (Figure 3F). Similarly, the nsP3 (2FA) mutant in WT U2OS cells showed a dramatically increased threshold concentration for condensation (Figure 3F).

To determine whether viral nsP3 protein could reach the concentration required for condensation during SFV infection, we estimated the cellular concentration of nsP3 at different time points post-infection with a standard curve created by recombinant gfp-nsP3 protein. We found that the nsP3 protein level could reach ∼3 μM at 4 hours post-infection (hpi) and ∼9 μM at 24 hpi (Figure S1M). The cellular G3BP1 protein level measured using the same method is ∼3 μM in 293T cells (Figure S1N), consistent with previous estimations^29^, thereby validating our method. Recombinant nsP3 alone can form condensates *in vitro*, with a threshold concentration between 15 to 20 μM, much higher than the threshold concentration measured in cells (Figures 3F and S2A). We found that G3BP1 addition, but not BSA control, can lower the threshold concentration of nsP3 for phase separation below 3 μM (Figure S2A). As a control, the enhancement effect was lost for nsP3(2FA) mutant protein, further demonstrating the critical role of G3BP1-nsP3 interaction in facilitating nsP3 condensation (Figure S2A). These results suggest that cellular nsP3 levels during viral infection can reach the condensation threshold in the early stage of infection.

Collectively, cellular and *in vitro* reconstitution data show the critical role of host G3BP1/2 proteins in facilitating viral nsP3 condensation.

### nsP3-G3BP co-condensate protects viral genomic RNA

To elucidate the function of nsP3 condensates in viral replication, we examined their spatial relationship with viral genomic RNA during infection. Using SFV-specific FISH probes, we detected a strong enrichment of viral RNA within nsP3 condensates (Figure 3G, top panel). Additionally, J2 antibody staining, which recognizes dsRNA intermediates during virus replication, revealed proximal localization to nsP3 condensates (Figure 3G, bottom panel). To determine whether nsP3 can bind viral RNA, we performed RNA-IP using gfp-nsP3 stable cells infected with SFV. A significant enrichment of viral RNA was observed in the nsP3 IP fraction (Figure 3H). Actin mRNA served as an internal control, showing no significant enrichment in the nsP3 IP fraction. Although nsP3 lacks canonical RNA-binding motifs, G3BP contains RRM and RGG motifs that facilitate RNA interaction. To assess whether the nsP3-G3BP interaction is required for nsP3-viral RNA binding, we transfected cells with the gfp-nsP3(2FA) construct and infected them with SFV. RNA-IP revealed that nsP3(2FA) abolished the interaction with viral RNA, suggesting that the interaction between nsP3 and viral RNA is likely mediated through G3BP (Figure 3H). A fluorescent polarization (FP) assay demonstrated that recombinant G3BP1 protein bound to the RNA probes with a K_D_ of approximately 0.5 μM, whereas nsP3 protein showed almost no RNA binding (Figure 3I). However, the addition of WT nsP3 enhanced the RNA binding of G3BP1 (Figure 3I). These results highlight the critical role of G3BP1 in mediating the nsP3-viral RNA interaction.

Based on the enrichment of virus RNA in nsP3 condensates, we hypothesize that these condensates could both protect the genomic RNA from host RNase and promote viral mRNA translation. We first demonstrated that multiple host RNases exhibit inhibitory effects on SFV in cells (Figure 3J). Using SAMHD1 and RNase L as examples, we showed that the viral inhibitory effect is highly dependent on the catalytic activity of both RNases (Figure 3K). Using an *in vitro* reconstitution system composed of nsP3, G3BP1, and a ∼1.4 kb *in vitro* transcribed SFV RNA fragment from nsP3, we found that the G3BP1-RNA condensates were efficiently disrupted by 1 ng/μl of RNase A addition (Figure 3L). However, the co-condensates of nsP3-G3BP1-RNA showed increased resistance to RNase A treatment within the same time window (Figure 3L). The addition of nsP3(2FA), which loses the direct interaction with G3BP, showed no protective effect. As a further control, the addition of Caprin1 to the G3BP-RNA system also showed no protective effect (Figure 3L). These results indicate a function for the viral nsP3 condensate in protecting viral RNA.

To test whether the condensation of nsP3 is essential for the protective effect, we further examined AUD deletion and nsP3 (3RE) mutants. We found that the protective effect was lost in both AUD deletion and 3RE mutants (Figure 3M). Furthermore, we found that both mutants exhibited increased dynamics *in vitro* compared to WT nsP3 (Figure S2B), suggesting that the material properties of condensates are critical for RNA protection. A previous report demonstrated that the liquid-like material properties of FUS low complexity domain (FUS_LCD) can be hardened by the G156E mutation^30^. We swapped the nsP3 AUD domain with WT and G156E mutant FUS-LCD. We found that both can rescue nsP3 condensation and FUS_LCD swap showed increased mobility compared to nsP3 (Figure S2C). The G156E swap exhibited reduced mobility compared to WT FUS_LCD. Consistently, FUS_LCD failed to protect RNA, whereas the G156E mutant swap showed a stronger protective effect compared to WT FUS_LCD (Figure S2D). In addition, we created a GST-FGDF fusion protein, which contains only the minimal region in nsP3 responsible for G3BP1 binding without the AUD domain. We observed an enhancement of G3BP1/RNA LLPS with GST-FGDF at a lower ratio; however, the GST-FGDF protein inhibited G3BP1 LLPS at a higher ratio (Figure S2E). FRAP analysis revealed that the GST-FGDF/G3BP1 condensates are more liquid-like compared to nsP3 condensates (Figure S2F), and no significant RNA protection effect was observed for the GST-FGDF/G3BP1 condensates (Figure S2G).

Collectively, these results indicate an intrinsic feature of the nsP3 AUD domain in maintaining gel-like state of the condensates and protecting RNA molecules inside condensates.

### nsP3 condensates promote protein translation

We have demonstrated the importance of nsP3 condensates in viral genome RNA stability. To further investigate whether the nsP3-G3BP condensates could directly regulate protein translation activity, we utilized the MCP-MS2 tethering system. We fused mCherry-G3BP1 with MCP and co-transfected it with a luciferase reporter that harbored a 12×MS2 repeat in the 3’UTR (Figure 4A). The MCP-mCherry-G3BP1 fusion significantly enhanced the translation activity of the luciferase reporter compared to the MCP-mCherry control or mCherry-G3BP1 alone (Figure 4B). To examine the role of nsP3 in regulating translation, we measured the luciferase activity in the presence of MCP-nsP3. We found that this construct also promoted the translation activity of the luciferase reporter (Figure 4C). To determine whether the translation-enhancing activity of nsP3 is dependent on G3BP1, we performed the assay in G3BP1/2 dKO cells and found that nsP3 could no longer enhance the translation activity (Figure 4C). The translation-enhancing activity of nsP3 was also abolished by the G3BP binding-deficient nsP3(2FA) mutant (Figure 4D), indicating that the translation enhancement by nsP3 is G3BP-dependent.

**Figure 4.**
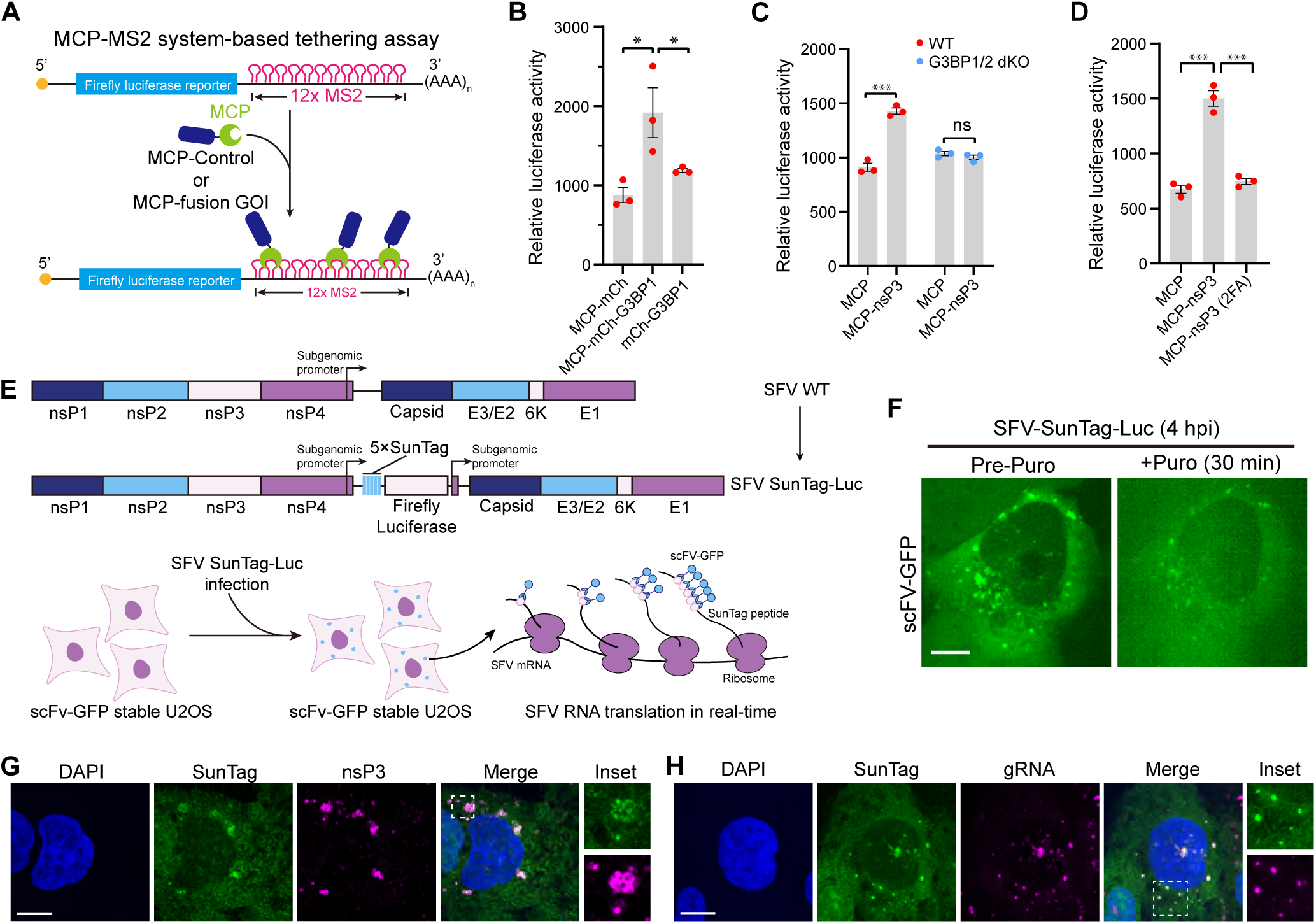
nsP3 condensates promote viral translation. (A) Diagram of the luciferase assay for protein translation activity. (B) Dual-luciferase activity measurement of MCP control, MCP-G3BP1, and untagged G3BP1 in HEK293T WT cells. Data are mean ± SEM with n=3 biological replicates. (C) Dual-luciferase activity measurement of MCP control, MCP-nsP3 in HEK293T WT and G3BP1/2 dKO cells. Data are mean ± SEM with n=3 biological replicates. (D) Dual-luciferase activity measurement of MCP control, MCP-nsP3 WT, and MCP-nsP3(2FA) mutant in HEK293T WT cells. Data are mean ± SEM with n=3 biological replicates. (E) Schematic of the recombinant SFV containing 5×SunTag arrays and a firefly luciferase CDS for viral translation imaging. (F) Representative images of SFV-SunTag-Luc translation treated with 100 μg/ml puromycin for 20 min. Scale bar, 10 μm. (G) Representative images of nsP3 antibody staining in scFv-GFP stable U2OS cells at 6 h after SFV-SunTag-Luc infection. Scale bar, 10 μm. (H) Representative images of vRNA FISH in scFv-GFP stable U2OS cells at 6 h after SFV-SunTag-Luc infection. Scale bar, 10 μm. Statistical significance in (B), (C), and (D) was assessed using an unpaired, two-tailed *t* test.

The enhancement of translation by nsP3 led us to hypothesize that nsP3 foci might serve as viral translation hubs during infection. To test this, we employed the SunTag reporter system, in which 24 repetitive copies of GCN4 epitopes (SunTag) were introduced at the 5’ end of a reporter gene. The sequence is recognized by GFP-fused single-chain antibody fragment (scFv-GFP) upon the emergence of the nascent SunTag peptide (Figure S3A). Under live cell imaging, scFv-GFP-bound SunTag-reporter appears as bright foci, indicating ongoing translation. We fused the SFV nsP4 CDS to the SunTag arrays as the reporter. With a dox-inducible promoter in the 5’ UTR and a 12×MS2 array in the 3’ UTR of the reporter, we can observe the inducible SunTag-nsP4-MS2 expression (Figure S3B). Treatment with puromycin to disassemble polysomes quickly abolished the GFP foci signal within 30 min, validating that the GFP foci reflect active translation (Figure S3C). To investigate the spatial relationship between active translation with nsP3 foci, we created MCP-nsP3 condensates capable of recruiting the reporter mRNA and nsP3 condensate without an MCP tag. Upon induction with doxycycline, we observed substantial GFP foci at the borders of nsP3 condensates in both conditions (Figures S3D and S3E). Live-cell imaging revealed the temporal co-movement of mCherry-nsP3 condensates with actively translating GFP foci, indicating that the GFP foci and nsP3 condensates interact during translation before eventually separating (Figure S3F). Collectively, these results suggest that nsP3 condensates can serve as active translation sites.

To determine the role of nsP3 condensates as translation sites during live virus infection, we inserted five copies of SunTag peptide arrays with firefly luciferase CDS into the spacer region between the SFV non-structural proteins and structural proteins (Figure 4E). The SunTag-luciferase insertion did not impair the viability of the recombinant SFV, as evidenced by the steady increase of luciferase activity over time upon viral infection (Figure S3G). Infection of scFv-GFP expressing cells showed GFP signal accumulation either as distinct spots or enclosed ring-like structures, both resembling the typical distribution patterns of nsP3 during viral infection (Figure S3H). The GFP foci were strongly diminished upon puromycin administration (Figure 4F), indicating sites of active translation. We observed good colocalization between these translation-active GFP foci and nsP3, as well as SFV genomic RNA (Figures 4G, 4H, and S3I). Together, these data suggest that nsP3-G3BP co-condensates can serve as active translation sites during viral infection.

### nsP3-G3BP1 interaction modulates SG dynamics

SGs perform antiviral functions during viral infection^31,32^. To investigate SG protein dynamics during SFV infection, we stained G3BP1 and eIF4G at various time points post-infection to visualize nsP3 condensates and SGs, respectively. At an early time point (6 hpi), all nsP3 foci were positive for G3BP1, and many cells also exhibited eIF4G-labeled SGs (Figure 5A, upper panel), with only a fraction of cells containing G3BP1-nsP3 condensates without the canonical SG marker eIF4G (Figure 5A, lower panel). However, the proportion of cells with eIF4G-positive SGs decreased over time, with less than 20% showing eIF4G-positive SGs at a later time point (24 hpi) (Figure 5B). We hypothesize that the elevated nsP3 protein levels during viral replication could alter the stoichiometric ratio between nsP3 and G3BP proteins, leading to a remodeled G3BP interaction network from SG formation to nsP3 viral condensate formation.

**Figure 5.**
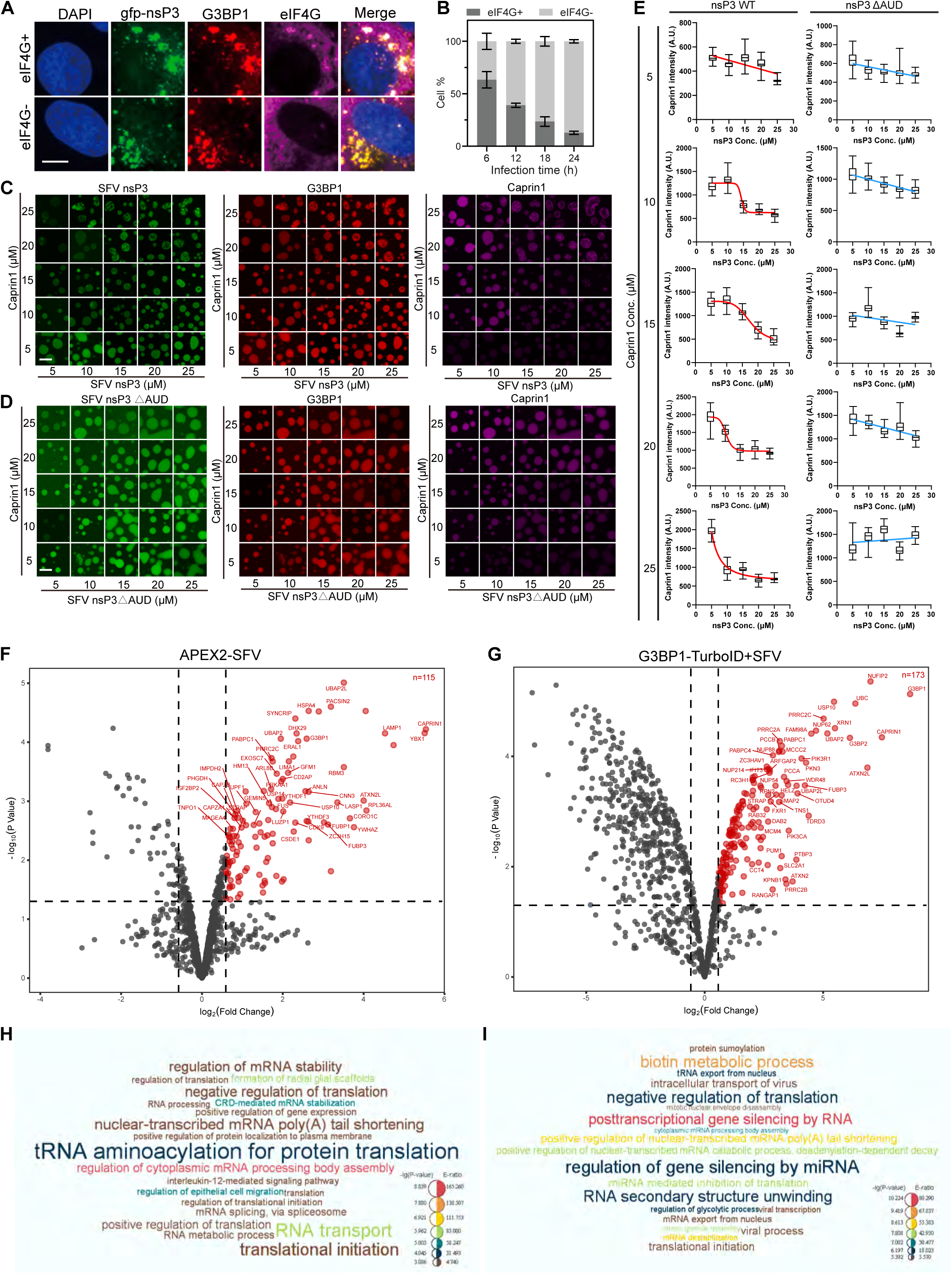
nsP3-G3BP1 interaction remodels antiviral stress granules. (A) Representative images of nsP3 and SG marker eIF4G in SG positive (top panel) and negative (bottom panel) cells. Scale bar, 10 μm. (B) Quantification of SG formation at different viral infection time points. Error bars indicate ± SEM; at each time point, n=3 measurement from 3 randomly selected fields of views (FOVs). (C) *In vitro* phase separation assay to assess the competition of Caprin1 by WT nsP3 protein. Scale bar, 5 μm. (D) *In vitro* phase separation assay to assess the competition of Caprin1 by nsP3 ΔAUD mutant protein. Scale bar, 5 μm. (E) Quantification of results in (C) and (D). Data are plotted as minimum to maximum; lines indicate the first quartile (lower), median, and third quartile (upper) with n=50 droplets for each box. For Caprin1 at 10, 15, 20, and 25 μM with nsP3 WT, a sigmoidal 4-parameter logistic curve (4PL) was used to fit the data; for Caprin1 at 5 μM with nsP3 WT and all Caprin1 with nsP3 ΔAUD, data were fitted with a simple linear regression model. (F and G) Volcano plots of APEX2-SFV mediated labeling of nsP3 condensate proteome (F) and TurboID-G3BP1 proteome with SFV infection (G). Red dots represent nsP3-interacted proteins with >1.5-fold enrichment compared to the control samples. See also Tables S1 and S2. (H and I) Top GO terms from (F) and (G).

To test this, we first compared the cellular concentrations of G3BP1/2 and nsP3 during virus infection. Our estimations, along with previous studies, showed that the combined concentration of G3BP1 and G3BP2 is approximately 2 μM in U2OS cells^25^ and 6 μM in 293T cells, respectively^29^ (Figure S1N). During viral infection, the nsP3 protein concentration can reach approximately 3 μM at 4 hpi (Figure S1M), and each nsP3 protein contains two G3BP binding motifs. This suggests that the nsP3 protein level at the early infection stage can efficiently compete with endogenous G3BP1 interactors, such as Caprin1, USP10, and UBAP2/2L. The stoichiometric ratio between nodes in an interacting network can modulate the dynamics of condensate^33–35^. Next, we tested the effect of competition between viral nsP3 and host interactors for G3BP1 binding on SG dynamics in cells by staining SG in the presence of nsP3 expression. Although many SG-localizing proteins are not components of nsP3 condensates (Figure 3A), they could re-localize to nsP3-G3BP1 co-condensates upon sodium arsenite (SA) treatment (Figure S4A). Our data are consistent with a recent report^36^ showing minimal impact of nsP3 expression on SG formation, in contrast to previous studies indicated that nsP3 inhibits SG formation through the hydrolase activity of the nsP3 Macro domain^37^ and the sequestration of G3BP proteins^38^. We further showed that nsP3 could form larger foci in response to SA in WT cells (Figure S4B). In WT U2OS cells, nsP3(2FA) mutant barely formed condensates (Figure 3E), but it could form condensates in response to SA treatment (Figure S4C). The threshold concentration for nsP3(2FA) forming condensate in cells was significantly reduced by SA treatment (Figure S4C). WT nsP3 rarely formed condensates in G3BP1/2 dKO cells but could form condensates at a much lower concentration in the presence of SA treatment (Figure S4D). These data suggest that nsP3 has an intrinsic condensation capacity in response to environmental stress. By disrupting nsP3-G3BP1 interaction, we found that nsP3(2FA) condensates are distinct from SG formation after SA treatment (Figure S4E). While examining cells with transiently transfected gfp-nsP3, we constantly observed two types of nsP3 granules: one larger and amorphous, and another smaller in size (Figures S4F and S4G). The SG marker Caprin1 was absent in smaller nsP3 foci, same as in nsP3 expressing cells (Figure 3A), which we referred to as nsP3 condensates; by contrast, Caprin1 was present in larger nsP3 foci, resembling canonical SGs. nsP3 also regulates SG fluidity, as revealed by the FRAP assay both *in vivo* and *in vitro* (Figures S4H and S4I), suggesting marked alteration of SG internal interaction network by nsP3. The evident differences in sizes and compositions indicate that SGs and nsP3 granules might function differently. Taken together, we conclude that viral nsP3 significantly rearranges the compositions and properties of SGs without an obvious effect on SG assembly.

Since nsP3 showed no obvious effect on SA-induced SG assembly, we then tested whether nsP3 could regulate SG disassembly. We found that nsP3 expression in cells led to lower enrichment of Caprin1 inside SGs (Figure S4J) and faster disassembly of SGs compared to control cells (Figure S4K). The effect was lost for the nsP3(2FA) mutant (Figure S4K), suggesting the importance of direct nsP3-G3BP1 interaction in regulating SG dynamics. We further tested the competition between SG and nsP3 condensate formation *in vitro*. Here, we used Caprin1, G3BP1, and RNA to reconstitute SGs and titrated increasing concentrations of nsP3 protein *in vitro* (Figure 5C). All three proteins can form co-condensates, and the titration of nsP3 efficiently excluded Caprin1 from the G3BP1-RNA condensates (Figures 5C, 5E, S5A, and S5B). We further demonstrated that condensation of nsP3 significantly affected Caprin1 exclusion, as the AUD deletion mutant showed a drastically reduced effect on Caprin1 exclusion at comparable concentrations (Figures 5D, 5E, S5A, and S5B). The influence of condensation was also shown with the nsP3 (2CA) and nsP3(3RE) mutants (Figures S5A and S5B). nsP3 contains two tandem FGDF motifs, which exhibit sub-micromolar binding affinity with G3BP1^39^. The binding affinity is comparable to that of the Caprin1 and USP10 binding to G3BP1^40^. To test the valence of FGDF on Caprin1 exclusion, we examined the effect of single F/A mutants (1FA). 1FA mutant also showed a reduced ability to exclude Caprin1 *in vitro* (Figures S5A and S5B). As a control, nsP3(2FA) mutant showed no effect on Caprin1 exclusion. The 1FA mutant retained the ability to co-phase separate with G3BP1 but with more spherical morphology for both 1FA mutants (Figure S5C). Furthermore, we found that WT nsP3 showed a dose-dependent effect on G3BP1 enrichment, and the effect was dramatically reduced for 1FA mutants (Figures S5D and S5E).

The above results suggest that nsP3 condensates modulate SG dynamics through direct competition with G3BP1 interactors.

### nsP3 remodels the G3BP1 interactome

To characterize the composition of the nsP3 condensates, we performed APEX2-mediated proximity labeling using the recombinant APEX2-SFV. Over 100 proteins were identified (Figure 5F), including previously known factors, such as G3BP1/2 and CD2AP^41^. In parallel, we generated a stable cell line expressing gfp-G3BP1-TurboID. The G3BP1 interactomes were mapped before (Figure S5F) and after SFV infection (Figure 5G). Both the nsP3 and G3BP1 interactomes were enriched with proteins involved in translation and mRNA metabolism pathways (Figures 5H, 5I, and S5G). Comparison between the nsP3 and SFV-infected G3BP1 interactomes identified many SG proteins, including G3BP1, Caprin1, and ATXN2L, suggesting a potential functional impact of the SG core protein network on SFV replication (Figure S5H). There is also a substantial change in the G3BP interactome following SFV infection. However, a significant portion of proteins are shared between the pre-and post-infection states, indicating a pre-existing G3BP complex that is remodeled upon infection (Figure S5I), similar to the remodeling observed after SG formation induced by sodium arsenite (SA) and heat shock^42,43^.

### Targeting nsP3 condensate inhibits viral replication

To test the effects of SG gene expression on virus replication, we expressed individual gene in cells and performed the SFV replication assay with the top hits from the APEX2-nsP3 MS (Figure 6A). A large number of SG proteins emerged as anti-SFV proteins, including Caprin1, UBAP2L, ATXN2/2L, YBX1, and USP10 (Figure 6A), which are key node proteins in the G3BP1 network for SG assembly^24^. Several major direct binders for G3BP1 stood out. Such G3BP1 direct binders (Caprin1, USP10, and UBAP2L) can directly interact with the NTF2L domain of G3BP1 via a conserved phenylalanine residue^42,44^. Based on the critical role of G3BP1 in mediating nsP3 condensate formation, we hypothesized that direct G3BP1 interactors could compete with nsP3 for G3BP1 binding (Figure 6B), thereby exerting a stronger antiviral effect. Via co-IP experiment, we found NUFIP2 as a new G3BP1 binder in addition to previously identified Caprin1, USP10, and UBAP2L (Figure S6A). However, ATXN2 and YBX1 showed no significant interactions (Figure S6A). Expression of NTF2L binding peptides from known G3BP1 interactors, such as USP10, Caprin1, UBAP2L, and SFV nsP3, can disrupt SG formation in cells^44^. We used this SG-disrupting assay to map the potential NTF2L binding peptide within NUFIP2. We identified the F511 residue in NUFIP2 as a critical residue mediating the G3BP1-NUFIP2 interaction (Figures S6B and S6C). Sequence alignment revealed the conservation of the phenylalanine residue among all NTF2L binders (Figure S6D). Through IP experiment coupled with AlphaFold3 prediction and computational docking, we confirmed the importance of NUFIP2 F511 residue in mediating the interaction with NTF2L domain (Figures S6E to S6G).

**Figure 6.**
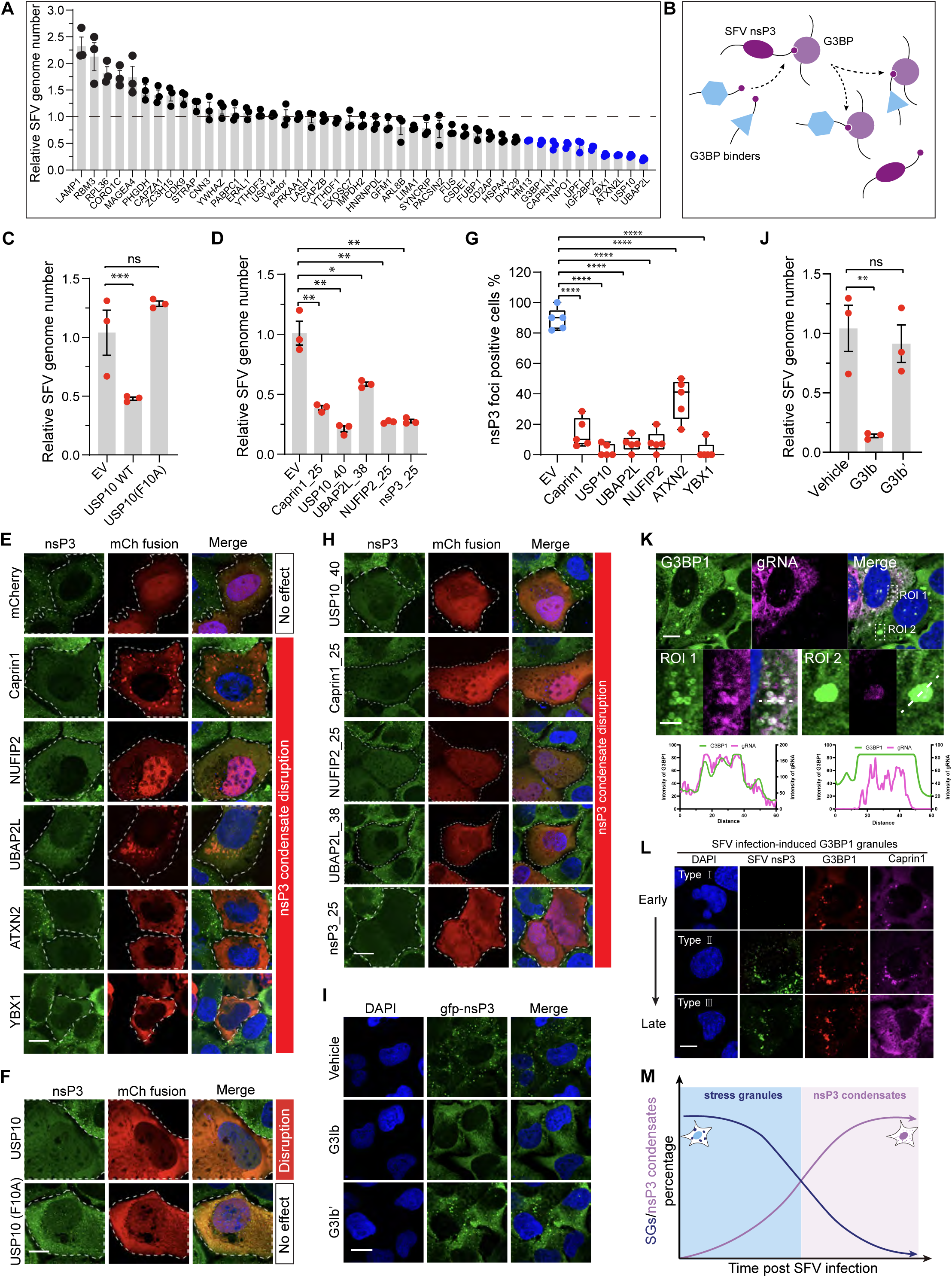
Targeting nsP3-G3BP interaction disrupts viral condensation and inhibits virus replication. (A) RT-qPCR results show the effect of gene overexpression on SFV replication in cells. Top hits are labeled in blue. Data are mean ± SEM with n=3 replicates. (B) Diagram of G3BP1 interactor remodeling during SFV infection. (C) RT-qPCR result of USP10 WT and its G3BP-binding deficient F10A mutant. Data are mean ± SEM with n=3 replicates. (D) RT-qPCR result of the antiviral effect from the indicated stress granule protein-derived short peptides. Data are mean ± SEM with n=3 replicates. (E) Representative images of nsP3 foci in the presence of the indicated SG proteins. Scale bar, 10 μm. (F) Representative images of nsP3 foci in the presence of USP10 WT and its F10A mutant. Scale bar, 10 μm. (G) Quantification of nsP3 positive cells co-expressed with the indicated SG components from (E) and (F). Data are plotted as a minimum to maximum; lines indicate the first quartile (lower), median, and third quartile (upper) with n=5 randomly selected FOVs. For each FOV, 10-15 cells were counted. (H) Representative images of nsP3 condensates in the presence of the indicated G3BP1-interacting peptides. Scale bar, 10 μm. (I) Representative images of nsP3 condensates upon G3Ib and G3Ib’ treatment. Scale bar, 10 μm. (J) Quantification of SFV genome in HEK293T cells treated with vehicle, G3Ib and G3Ib’. Data are mean ± SEM with n=3 replicates. (K) Representative immunofluorescence (IF) and FISH images showing the colocalization of viral genome RNA and G3BP1 positive stress granules. Scale bars, 10 μm for the upper panel and 200 nm for the inset. (L) Representative images showing the patterns of nsP3 structures and Caprin1 labeled SG at different stages of SFV infection. Scale bar, 10 μm. (M) A model showing the competition of nsP3 condensate and SG determined by nsP3-G3BP1 stoichiometry. Statistical significance in (C), (D), (G), and (J) was assessed using an unpaired, two-tailed *t* test.

Having shown the common G3BP1 interacting mechanism, we next asked whether the SFV inhibiting effect relies on G3BP1 interaction. We found that USP10 can inhibit SFV replication (Figures 6A and 6C), but the G3BP1 interaction-disrupting mutant, F10A, failed to inhibit SFV (Figure 6C). In addition, G3BP1 interacting peptides from USP10 and other binders are all sufficient to inhibit SFV (Figure 6D). Due to the significant role of nsP3 condensates in promoting SFV replication in cells, we investigated the effect of SG proteins and peptides on nsP3 condensate formation. We observed a broad inhibiting effect on viral condensates for the tested SG proteins, including Caprin1, NUFIP2, UBAP2L, ATXN2, and YBX1 (Figures 6E and 6G). USP10 can also disrupt nsP3 condensates, and the F10A mutation abolished the effect (Figures 6F and 6G). As expected, the five NTF2L binding peptides were sufficient to disrupt nsP3 condensates as well (Figure 6H). A chemical compound, G3Ib, was developed based on nsP3 FGDF peptide^45^, and we found it potently disrupted nsP3 condensate formation, whereas the inactive enantiomer, G3Ib’, was ineffective (Figure 6I). The compound also demonstrated a strong inhibitory effect on SFV replication (Figure 6J). To support the cellular effect, we further showed that G3Ib reduced the nsP3-G3BP1 interaction (Figure S6H) and co-phase separation of nsP3-G3BP1 *in vitro* (Figure S6I).

Full-length G3BP1 also exhibited antiviral effects upon overexpression as demonstrated in the virus-inhibiting screen shown in Figure 6A. We observed a substantial fraction of cells expressing G3BP1 exhibiting spontaneous stress granules (sSG) (Figure 6K). In sSG-containing cells, viral genomic RNA levels were significantly reduced, and there was an enrichment of viral RNA within SGs (Figure 6K). The formation of sSGs correlated with a broad translation shutdown as revealed by the puromycin incorporation assay, in which all tested SG-promoting proteins inhibited translation (Figure S6J). The role of sSG formation in bulk translation shutdown was further confirmed by examining G3BP1 mutants deficient in sSG formation, which failed to induce translation shutdown (Figure S6K). These results suggest that the condensation of mRNA within SGs can suppress its translation activity. This is further supported by the *in vitro* translation assay, whereby the translation activity of the firefly luciferase mRNA was diminished upon G3BP1 condensation (Figure S6L). Therefore, despite G3BP1 overexpression leading to colocalization with nsP3, its ability to promote SG formation and condense viral RNA contributes to its antiviral effects. Consistently, we observed a gradual shift of SG formation (marked as Caprin1+; G3BP1+) to nsP3 condensate formation (marked as Caprin1-; G3BP1+) from early to late stages of virus infection (Figure 6L). Using puromycin incorporation assay, we found a significant restoration of translation activity in nsP3-positive cells that were negative for SG marker Caprin1 (Figure S6M). Additionally, there was an increase in lower molecular weight protein translation during the late stages of SFV infection, which corresponded well to the increased translation of viral proteins (Figure S6N). Using SunTag reporter cells, we found that SunTag foci were eliminated in the presence of sSG induced by G3BP1 overexpression, in contrast to the active translation of nsP3 condensate (Figure S6O). Consistently, global translation was arrested in cells with larger sSG but de-repressed in cells with smaller canonical nsP3 condensates as revealed by the puromycin incorporation assay (Figure S6P). Consistent with the idea that SGs inhibit the translation of sequestered mRNAs, we did not observe any translating SunTag foci in the sSG-containing cells during SFV-SunTag infection (Figure S6Q), indicating the opposite effects of SGs and nsP3 condensates on viral mRNA translation.

Thus, anti-viral SGs and pro-viral nsP3 condensates are competing condensates centered on the G3BP protein node (Figure 6M), providing a rationale for targeting the nsP3-G3BP1 interaction as an antiviral intervention strategy.

### nsP3 condensation is conserved in alphavirus family

To assess the prevalence of nsP3 condensation and G3BP interaction among alphaviruses, we aligned the nsP3 protein sequences from the 32 currently identified alphaviruses (Figure S7A). Traditionally, the alphavirus family has been divided into two geographical sub-classes: New World and Old World alphaviruses^15^. This division aligns well with phylogenetic tree analysis based on nsP3 amino acid sequences (Figure S7A). The modular structure of nsP3 is conserved, with the C-terminal hypervariable domain (HVD) exhibiting the most variation. Alignment analysis revealed the conservation of several critical regions involved in G3BP1 binding. Among the 32 members, 17 contain the “FGDF”-like motif (Figures S7A and S7B), predominantly in the Old World family. This analysis suggested that the dependency on G3BP1 is likely a common feature of the Old World alphaviruses. The conservation of the tandem “FGDF” pattern also supports our finding on the importance of FGDF valence in regulating nsP3 condensate and SG dynamics. Genetic studies have demonstrated the essential role of this tandem motif in virus fitness; removing one FGDF by nsP3 C-terminal truncation or introducing F/A mutation to block G3BP interaction resulted in approximately a 100-fold reduction in SFV virus titer^28,39^.

Many New World alphaviruses, including VEEV, lack the FGDF motif for G3BP binding but instead utilize FXR1/FXR2/FMR1 (FXR) proteins for optimal replication in human cells^46^. Previous reports have shown that two repetitive FXR-binding motifs are present within VEEV (TC83 strain) nsP3 HVD^47^. To identify the key residues for FXR binding, we generated nine mutants with alanine substitution (Figure S7C). IP-WB analysis showed that FXR association was specifically lost with the VIT residues mutation (Figures S7C and S7D). IF experiment revealed that the recruitment of FXR1 into the VEEV nsP3 foci was abolished in the VIT mutant (Figure S7E). Other alphaviruses within the New World also harbor a degenerative VIT motif (Figures S7A and S7F), suggesting that the VIT motif, particularly the isoleucine residue, might be critical for FXR protein interaction.

Given their epidemic potential and distinct phylogenetic differences compared to SFV, we specifically examined the nsP3 from EEEV, WEEV, and VEEV. Expression of each nsP3 protein in U2OS cells resulted in distinct cytoplasmic foci (Figures 7A to 7C). FRAP and live cell imaging demonstrated that these three nsP3 can form condensates when expressed in cells (Figures 7A and 7B). All of these nsP3 showed a slow recovery in FRAP assay, indicating gel-like features of the nsP3 condensates. The two cysteines are conserved among nsP3 proteins, and 2CA mutants of EEEV, WEEV, and VEEV also disrupted condensate formation in cells (Figure 7C). The IP experiment showed a more robust interaction with G3BP1 for SFV nsP3 compared to the other three nsP3 analyzed. Conversely, the interaction with FMR1 was stronger for the New World alphaviral nsP3 compared to Old World SFV (Figure 7D). Although there is no canonical “FGDF” binding motif in EEEV/WEEV nsP3s, they can still interact with G3BP1, albeit with weaker affinity compared to SFV nsP3. We found that EEEV, WEEV, and VEEV nsP3 could still form condensate in G3BP1/2 dKO cells (Figure 7E), indicating that the condensation of New World nsP3 in cells does not rely on G3BP proteins. The higher sensitivity of SFV to G3BP1/2 KO is likely due to the critical dependence of SFV nsP3 condensation on G3BP1/2 proteins compared to other nsP3 (Figures 7D and 7E). Consistent with the IP results, endogenous G3BP1 was recruited into SFV/EEEV/WEEV nsP3 condensates but not into the VEEV nsP3 condensates (Figure 7F). Under SA treatment, canonical SGs could still form and co-localize with SFV/EEEV/WEEV nsP3 condensates but were separated from VEEV nsP3 condensates (Figure 7G). In contrast, FMR1 was recruited into EEEV/WEEV/VEEV nsP3 condensates but not into SFV nsP3 condensates (Figure 7H).

**Figure 7.**
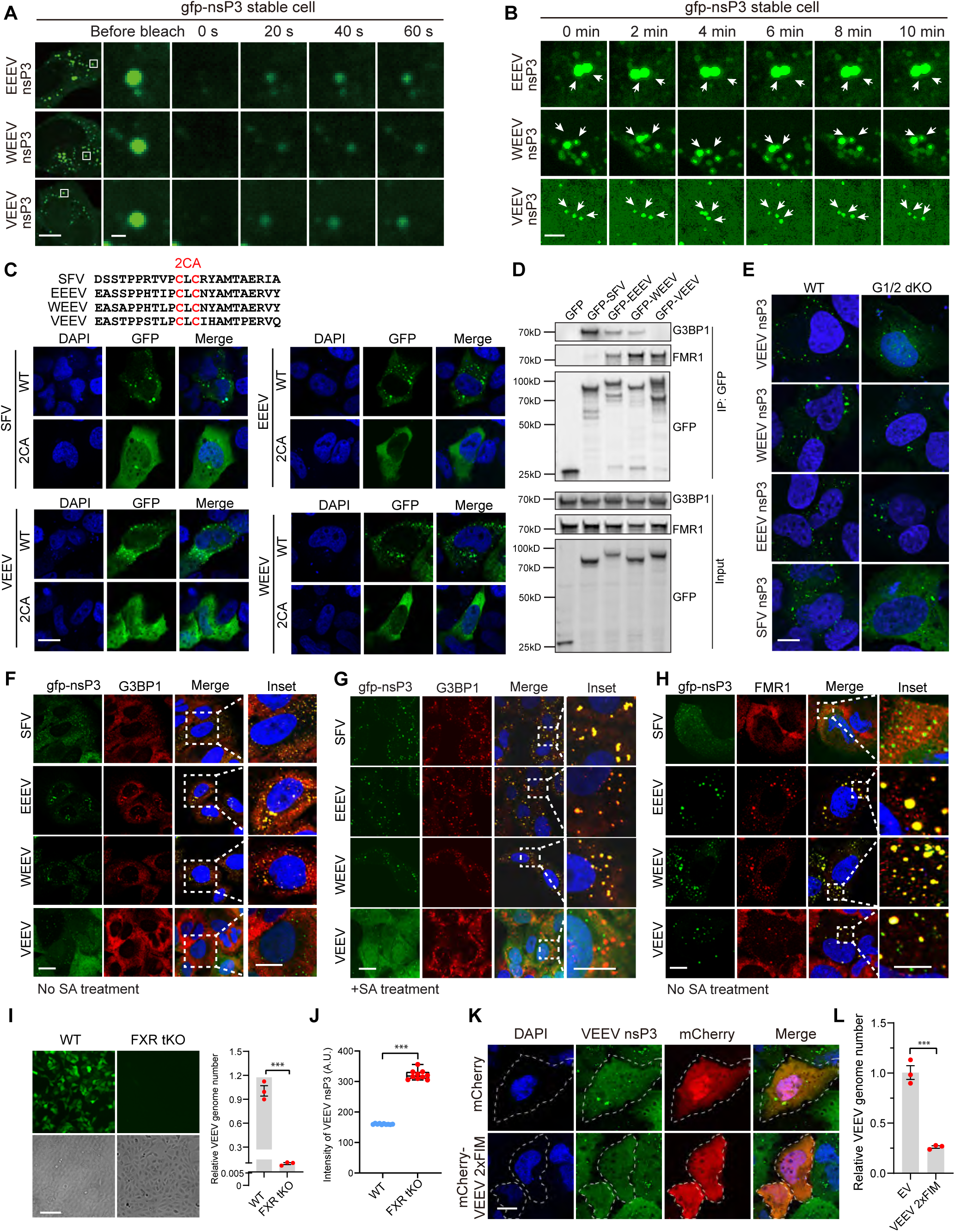
nsP3 condensation is conserved in alphavirus. (A) Representative images of FRAP experiments on EEEV/WEEV/VEEV nsP3 proteins in cells. Scale bar, 10 μm and 1 μm for the inset. (B) Representative images of slow fusion of EEEV/WEEV/VEEV nsP3 droplets in cells. Scale bar, 5 μm. (C) Representative images of WT and 2CA mutants of each nsP3 protein in cells. The alignment of the two cysteines is shown on the top. Scale bar, 10 μm. (D) Representative image of IP-WB to show the interaction between the four nsP3 proteins and G3BP1/FMR1. nsP3 construct was transfected into 293T cells, and WB was performed with G3BP1 or FMR1 antibodies. (E) Representative images of SFV/WEEV/EEEV/VEEV nsP3 protein in WT and G3BP1/2 dKO cells. Scale bar, 10 μm. (F) Representative images of localization between nsP3 and G3BP1. Scale bars, 10 μm and 5 μm for the inset. (G) Representative images of localization between nsP3 and G3BP1 with SA treatment. Scale bars, 10 μm and 5 μm for the inset. (H) Representative images of localization between nsP3 and FMR1. Scale bars, 10 μm and 5 μm for the inset. (I) Left: representative images of VEEV(GFP) replication in WT and FXR1/FXR2/FMR1 triple KO (FXR tKO) U2OS cells. The top row shows the fluorescent signal from VEEV(GFP), and the bright field below indicates the cell density. Scale bar, 10 μm. Right: RT-qPCR quantification of VEEV genome in U2OS WT and FXR tKO cells. Data are mean ± SEM with n=3 replicates. (J) Quantification of relative threshold fluorescence for VEEV nsP3. Data are plotted as a minimum to maximum; lines indicate the first quartile (lower), median, and third quartile (upper). n=10 values in the determined thresholding bin. (K) Disruption of VEEV nsP3 condensate formation by FXR-interacting motif peptide from VEEV (VEEV 2×FIM). Scale bar, 10 μm. (L) RT-qPCR analysis of the VEEV replication in HEK293T transfected with the FXR binding peptide VEEV 2×FIM. Data are mean ± SEM with n=3 replicates. Statistical significance in (I), (J), and (L) was assessed using an unpaired, two-tailed *t* test.

To test whether the VEEV nsP3-FMR1 interaction follows a similar principle to the SFV nsP3-G3BP1 interaction in virus replication, we created FXR1/FXR2/FMR1 triple KO U2OS cells. We successfully recapitulated the requirement of these three paralogs for GFP-VEEV (TC83) replication (Figure 7I). RT-qPCR showed a >100-fold reduction of VEEV RNA levels in FXR tKO cells (Figure 7I). The threshold concentration for VEEV nsP3 was dramatically elevated in FXR tKO cells (Figure 7J). Based on the FXR1 binding motif inside VEEV nsP3^46^ (Figures S7A and S7F), we generated a peptide to inhibit VEEV nsP3 condensation in cells (Figure 7K). Expression of this peptide exhibited antiviral effects against VEEV, suggesting that the nsP3-FXR interaction may be targetable for the New World alphaviruses (Figure 7L). EEEV and WEEV nsP3 could colocalize with both G3BP and FXR family proteins (Figures 7F to 7H). Therefore, we co-transfected peptides targeting both G3BP and FXR and observed the disruption of nsP3 condensates from both EEEV and WEEV (Figure S7G), indicating that combinatorial treatments could be feasible.

The New World alphaviruses can either utilize FXRs specifically or in combination with G3BPs for viral replication^46^. Thus, FXRs may function analogously to G3BPs in regulating viral and host translation. Using the MCP-MS2 tethering assay, we demonstrated that MCP-EEEV/WEEV/VEEV nsP3 could enhance tethered luciferase translation as well (Figure S7H). The enhancement of reporter translation by VEEV nsP3 is abolished in FXR1/FXR2/FMR1 tKO cell (Figure S7I). Collectively, these data suggest that co-condensation between alphaviral nsP3 and host translation-enhancing RBPs is likely a conserved mechanism and it is feasible to target alphavirus by disrupting viral nsP3 condensates.

## Discussion

Our study using the alphavirus SFV as a model demonstrated the intrinsic condensation capability of the nsP3 protein and its ability to form co-condensate with host RBPs. These features are conserved across alphavirus family members. nsP3 can interact with host RBPs, such as G3BP1/2, FXR1/FXR2/FMR1, to facilitate viral RNA binding, thereby increasing genome stability and promoting translation efficiency. Nascent nsP3 protein can form a complex with host G3BP protein due to its high binding affinity and valency upon initial release from viral positive-strand RNA template. G3BP has been shown to enhance the stability and promote the translation of target RNA^48^. Using *in vitro* reconstitution and cellular infection assays, our work demonstrated that the condensation of nsP3/G3BP1 can protect viral RNA from host RNase degradation, enhance viral RNA stability, and promote viral RNA translation efficiency. The viral nsP3 exhibits intrinsic features in condensation, which can be facilitated by host RBPs such as G3BP and FXR proteins.

Protein condensation can be initiated through interactions between intrinsically disordered regions (IDRs) or structured domains capable of providing multivalent binding. Although nsP3 contains a C-terminal HVD as an IDR, the primary determinant of condensation lies within the highly conserved and structured AUD domain. We showed that critical cysteine residues in the AUD domain are essential for nsP3 condensation and that nsP3 condensates are necessary for optimal viral replication. The essential role of AUD in nsP3 condensation is supported by the recent Cryo-EM structure of Chikungunya virus (CHIKV) nsP3-AUD^22^. Given the high similarity of AUD across alphaviral members, nsP3 condensation may be a conserved mechanism in alphaviruses for organizing intracellular replication complexes.

Stress granule assembly correlates with translational shut-off, which can suppress viral protein translation and is thus regarded as a general antiviral mechanism. SGs exert antiviral functions through several mechanisms, including direct condensation with viral RNA and activation of immune signaling pathways^31,32^. Many viral proteins inhibit SG assembly through either cleavage of G3BP or sequestration of G3BP from other interconnected SG proteins^49^. We demonstrated that nsP3 condensation contributes to host SG disassembly by modulating the composition of the G3BP interaction network. Our findings and a recent study from Dr. Ilya Frolov^50^ suggested novel mechanisms for how alphaviral nsP3 may modulate SG dynamics in addition to the macrodomain-mediated mechanism^37^. Therefore, the formation of SGs and viral nsP3 condensates represents a tug-of-war centered on G3BP proteins (Figure S7J).

The essentiality of G3BP1 in SFV replication and the high mutability of the HVD region of nsP3 suggest that the binding of novel host RBPs may be an evolvable trait for the alphavirus family. The identification of the New World alphavirus reveals the evolution of novel host RBP binding. Thus, a similar principle likely applies to the New World alphavirus. For instance, the valence of nsP3-FXR1 family RBP interaction is not fixed in several pandemic VEEV stains. The disease severity caused by any VEEV strain is likely correlated with the FXR binding valence^47,51^, suggesting that the binding specificity and valence of host RBPs are ongoing evolvable traits for many New World alphaviruses.

Finally, by targeting the nsP3-RBP interaction with NTF2L binding peptide and a derivative compound, we demonstrated the proof of principle in modulating viral replication in cells. The conservation of nsP3-RBP interactions in alphavirus and the conservation of G3BP as a host factor targeted by diverse viruses^52–55^ suggest that broad-spectrum antiviral agents could be developed based on intracellular viral replication steps, which are critically conserved during viral replication.

## Supporting information

List of human proteins identified through APEX2-mediated proximity labeling for SFV nsP3 protein

List of human proteins identified through TurboID-mediated proximity labeling for G3BP1 protein with SFV infection

List of human proteins identified through TurboID-mediated proximity labeling for G3BP1 protein without SFV infection

FISH probes used for SFV genomic RNA detection

qPCR primers used in this study

## Acknowledgement

We are grateful for the gift of the plasmid encoding the Semliki Forest Virus (SFV) from Dr. Andres Merits at University of Tartu, Estonia. We thank Dr. Weirui Ma, Zhejiang University, China, for the SunTag plasmid system and technical guidance. We thank the technical support from the Westlake Biomedical Research Core Facility. This work was supported by grants from the National Key Research and Development Project of China (2025YFC3409700 to P.Y.), the National Natural Science Foundation of China (32470733 and 32170696 to P.Y.), the Center of Synthetic Biology and Integrated Bioengineering of Westlake University (WU2022A002), “Pioneer” and “Leading Goose” R&D Program of Zhejiang (2024SSYS0030), and the HRHI program of Westlake Laboratory of Life Sciences and Biomedicine (W101110536022201).

## Author contribution

Y.L. performed cellular experiments, viral infection, and *in vitro* reconstitution experiments. Y.L., Z.Y., Q.C., X.C., Y.Z., Z.Z., L.T., Z.W., Y.M., and W.F. participated in plasmid construction, cell line generation, and microscopy studies. Y.H. and Y.Z. participated in the super-resolution microscope study. J.D. and J.H. participated in the molecular docking study. Z.Y., J.W., H.K., and J.P.T participated in the G3BP1 targeting compound study. P.Y. conceived and supervised the project with input from J.P.T.. P.Y. wrote the manuscript with input from all authors.

## Supplemental information

Document S1. Figures S1–S7

Table S1. List of human proteins identified through APEX2-mediated proximity labeling for SFV nsP3 protein.

Table S2. List of human proteins identified through TurboID-mediated proximity labeling for G3BP1 protein with SFV infection.

Table S3. List of human proteins identified through TurboID-mediated proximity labeling for G3BP1 protein without SFV infection.

Table S4. FISH probes used for SFV genomic RNA detection. Table S5. qPCR primers used in this study.

**Figure S1. SFV nsP3 forms condensates. Related to Figure 1 and Figure 2.**

(A) Diagram of SFV encoded proteins.

(B) PONDR analysis of four SFV non-structural proteins (nsP).

(C) Representative images of each nsP expression in cells. Scale bar, 10 μm.

(D) Representative images of WT and transmembrane (TM) domain deletion mutant of nsP1 in cells. Scale bar, 10 μm.

(E) Representative images of nsP3 droplet fusion. Scale bar, 10 μm.

(F) Images for nsP3 antibody verification. Scale bar, 10 μm.

(G) Representative images of nsP3 droplet fusion during SFV infection. Scale bar, 2 μm.

(H) Images for staining of APEX2-SFV infected cells. Scale bar, 10 μm.

(I) Alignment between SFV_AUD and CHIKV_AUD. The two cysteines and three arginine residues are underlined by the asterisk.

(J) Salt concentration-dependent phase separation of nsP3 *in vitro*. Scale bar, 20 μm.

(K) IP-WB to map the nsP3 domains required for self-interaction.

(L) RT-qPCR results showing the relative expression level of SFV receptors in WT and G3BP1/2 dKO cells. Data are mean ± SEM with n=3 replicates.

(M) Quantification of nsP3 protein level during viral infection.

(N) Quantification of G3BP1 protein level in 293T cells. The volume of 293T cells is 1×10^-12^ L^56^.

Statistical significance in (L) was assessed using an unpaired, two-tailed *t* test.

**Figure S2. Gel-like nsP3 condensates shield viral RNA. Related to Figure 3.**

(A) Representative images of co-phase separation assay of nsP3-G3BP1 *in vitro*. BSA and the nsP3(2FA) mutant were used as controls. Scale bar, 20 μm.

(B) FRAP assay showing the dynamic of WT and AUD deletion mutant of nsP3 condensates *in vitro*. A quantification curve is shown on the right. Data are mean ± SEM with n=10 droplets measured. Scale bar, 5 μm.

(C) FRAP assay showing the dynamics of nsP3_FUS_LCD WT and nsP3_FUS_LCD (G156E) mutant dynamic of nsP3 condensates *in vitro*. A quantification curve is shown on the right. Data are mean ± SEM with n=5 droplets measured. Scale bar, 5 μm.

(D) RNA protection assay with WT and nsP3 swap mutants. RNA labeled with Cy5 was used for imaging. Scale bar, 5 μm.

(E) Phase separation of GST-FGDF with G3BP1 *in vitro*. Scale bar, 20 μm.

(F) FRAP assay showing the dynamic of GST-FGDF proteins compared to nsP3 *in vitro*. Data are mean ± SEM with n=10 droplets measured. Scale bar, 5 μm.

(G) Representative images of RNA protection effect of GST-FGDF. Scale bar, 20 μm. Statistical significance in (B), (C), and (F) was assessed using a two-way ANOVA, Šídák’s multiple comparisons test.

**Figure S3. nsP3 condensates show protein translation activity. Related to Figure 4.**

(A) Workflow of creating the SunTag-nsP4-MS2 reporter U2OS cell line for SunTag imaging.

(B) Representative images of SunTag-nsP4 translation upon doxycycline treatment for 2 hours. Scale bar, 20 μm.

(C) Representative time-lapse images of SunTag-nsP4 translation treated with 100 μg/ml puromycin. Scale bar, 10 μm.

(D and E) Representative images of SunTag-nsP4 translation in cells co-expressed with MCP-mCherry-nsP3 (D) and MCP-free mCherry-nsP3 (E). Scale bar, 5 μm and 1 μm for the inset.

(F) Spatiotemporal tracking of SunTag-nsP4 translation signal (arrow) and an MCP-free mCerry-nsP3 condensate. Scale bar, 1 μm.

(G) Measurement of luciferase activity in U2OS cells infected with the SFV-SunTag-Luc for the indicated time.

(H) Representative images of scFv-GFP stable U2OS cells at 6 h after SFV-SunTag-Luc infection. Two typical GFP foci patterns were demonstrated. Scale bar, 10 μm and 2 μm for the inset.

(I) Time lapse imaging and single particle tracking of SFV-SunTag-Luc translation signal and nsP3 condensates in scFv-GFP/mCherry-nsP3 double stable U2OS cells at 6 h after SFV-SunTag-Luc infection. Arrows indicated the merged SunTag foci and nsP3 foci. Scale bar, 1 μm.

**Figure S4. Inter-relationship and mutual regulation between nsP3 condensates and SG. Related to Figure 5.**

(A) gfp-nsP3 expressing cells were treated with sodium arsenite (SA) and stained with SG markers, including G3BP1/2, TIA1, Caprin1, PABP1, eIF4G, and eIF3η. Scale bar, 10 μm.

(B) Representative images of gfp-nsP3 with and without SA treatment. Scale bar, 10 μm.

(C) Representative images of gfp-nsP3(2FA) with and without SA treatment in WT U2OS cells. Below is the quantification of threshold fluorescence for gfp-nsP3(2FA). Data are plotted as a minimum to maximum; lines indicate the first quartile (lower), median, and third quartile (upper). n=10 values in the determined thresholding bin. See Methods for defining the bin. Scale bar, 10 μm.

(D) Representative images of gfp-nsP3 WT with and without SA treatment in G3BP1/2 dKO cells. Below is the quantification of threshold fluorescence for gfp-nsP3 WT. Data are plotted as a minimum to maximum; lines indicate the first quartile (lower), median, and third quartile (upper). n=10 values in the determined thresholding bin. See Methods for defining the bin. Scale bar, 10 μm.

(E) gfp-nsP3(2FA) expressing cells were treated with sodium arsenite (SA) and stained with SG markers to compare with images in (A). Scale bar, 10 μm.

(F) Representative immunofluorescence images of gfp-nsP3 transfected U2OS cells co-stained with G3BP1 and Caprin1. Scale bar, 20 μm.

(G) Calculation of the relative nsP3-G3BP1 foci area, n=50.

(H) *In vitro* FRAP curve of nsP3 RNA-induced G3BP1 droplets (n=10) in the presence or absence of 25 μM nsP3.

(I) *In vivo* FRAP curve of G3BP1 in mCherry-G3BP1 stable U2OS cells co-expressed with gfp or gfp-nsP3 and treated with SA for 1 hour, n=5.

(J) Localization of Caprin1 inside SG is reduced in the presence of nsP3. Control and nsP3-expressing cells were treated with SA and stained for Caprin1 and G3BP1. Scale bar, 10 μm.

(K) Cells were collected at the indicated time points after SA washout and subject to immunofluorescence for G3BP1 and Caprin1. The ratio of SG-positive cells (G3BP1^+^/Caprin1^+^) was quantified. Error bars indicate ± SEM. n=3 FOVs with at least 20 cells counted for each FOV.

Statistical significance in (C), (D), and (G) was assessed using an unpaired, two-tailed *t* test. Statistical significance in (H) and (I) was assessed using a two-way ANOVA, Šídák’s multiple comparisons test.

**Figure S5. Valence-dependent competition of Caprin1 in nsP3 condensates and remodeling of G3BP1 interactome upon SFV infection. Related to Figure 5.**

(A) *In vitro* phase separation assay to assess the competition of Caprin1 by WT and mutant nsP3 protein. Scale bar, 20 μm.

(B) Quantification of results in (A). Data are plotted as minimum to maximum; lines indicate the first quartile (lower), median, and third quartile (upper) with n=30 droplets for each box.

(C) Representative images of nsP3-G3BP1 phase separation in the presence of RNA. The nsP3 WT, F451A, and F468A mutants were compared. Scale bar, 20 μm.

(D) Fluorescent channel to show the enrichment of G3BP1 protein inside nsP3-G3BP1 condensates. Scale bar, 20 μm.

(E) Quantification of G3BP1 enrichment in (D) as measured as partition coefficient value. Data are plotted as a minimum to maximum; lines indicate the first quartile (lower), median, and third quartile (upper) with n=10 droplets for each box.

(F) Volcano plot of TurboID-G3BP1 interactome without SFV infection. Red dots represent G3BP1-interacted proteins with >1.5-fold enrichment compared to the control samples. See also Table S3.

(G) Top GO terms from G3BP1 interactome without virus infection.

(H) Comparative analysis of APEX2-SFV labeled nsP3 proteome and G3BP1 proteome.

(I) Comparative analysis of TurboID-G3BP1 labeled proteome before and after SFV infection.

**Figure S6. Investigation of G3BP1 binders and the effect of SG on translation activity. Related to Figure 6.**

(A) IP-WB to show the interaction between SG proteins with G3BP1 in cells.

(B) Representative images of the expression of NUFIP2 full-length and fragments on SG formation. The fragment inhibiting SG is highlighted in red box. Scale bar, 10 μm.

(C) Representative images of the expression of NUFIP2 F/A mutants on SG formation. The mutant losing SG-inhibiting ability is highlighted in red box. Scale bar, 10 μm.

(D) Alignment showing the critical phenylalanine residue in NTF2L direct binders.

(E) IP-WB to show the interaction of NTF2L binding peptides with G3BP1 in cells.

(F) AlphaFold3 prediction of NUFIP2 binding with NTF2L and their alignment with USP10, Caprin1, and nsP3 peptides. Structures of G3BP1 NTF2L in complex with the indicated peptides are shown. PDB IDs are indicated for each sequence. The side chain of relevant phenylalanine residues is shown. Corresponding residues in the sequence are labeled in red.

(G) Overlay of the conformations of the four NTF2L binding peptides to show the conserved orientation of the critical phenylalanine in NTF2L binding.

(H) IP-WB to show the dose-dependent disruption of nsP3-G3BP1 interaction by G3Ib compound. G3Ib’ is used as control.

(I) Representative images to show the dose-dependent disruption of nsP3-G3BP1phase separation *in vitro*. Scale bar, 10 μm.

(J) Puromycin incorporation assay to detect protein translation activity in cells expressing individual SG protein. Scale bar, 10 μm.

(K) Puromycin incorporation assay to detect protein translation activity with G3BP1 mutant expression. Scale bar, 10 μm.

(L) Top: G3BP1 phase separates with RNA *in vitro*. Bottom: *In vitro* translation activity assayed with increasing concentrations of G3BP1. Data are mean ± SEM with n=3 replicates. Scale bar, 10 μm.

(M) Puromycin incorporation assay to detect protein translation activity of cells infected with SFV. Quantification of puromycin signal inside Caprin1 positive and negative cells is shown on the right. Data are plotted as a minimum to maximum; lines indicate the first quartile (lower), median, and third quartile (upper) with n=7 FOVs in which 10-15 cells were counted for each FOV. Scale bar, 10 μm.

(N) WB of cellular global translation activity upon SFV infection at the indicated time points.

(O) Representative images of SunTag-nsP4 translation in cells co-expressed with mCherry, mCherry-G3BP1 and mCherry-SFV nsP3. Scale bar, 10 μm.

(P) Immunofluorescence of puromycin in U2OS cells co-transfected with gfp-nsP3 and mCherry-G3BP1. The representative images of cells containing canonical SGs (upper) or typical nsP3 condensates (lower) were shown. Scale bar, 10 μm.

(Q) Representative images of SG-containing scFv-GFP/mCherry-G3BP1 stable cells showing the absence of GFP-SunTag foci at 6 h after SFV-SunTag-Luc infection. Scale bar, 5 μm and 2 μm for the inset.

**Figure S7. Conservation of alphaviral nsP3 proteins. Related to Figure 7.**

(A) Phylogenetic tree of alphavirus family based on nsP3 protein sequence.

(B) Alignment of 17 Old World alphavirus family members encoding nsP3 proteins with conserved G3BP binding motifs.

(C) Domain diagrams of WT and VEEV nsP3 mutants. Alanine mutations were indicated by short lines for each mutant.

(D) Immunoprecipitation of transfected gfp-VEEV nsP3 mutants and endogenous FXR1 in U2OS cells.

(E) Immunofluorescence of endogenous FXR1 in U2OS co-transfected with gfp-VEEV nsP3 mutants. Mutants losing FXR1 recruitment is highlighted in red box. Scale bar, 10 μm.

(F) Alignment of FXR binding motifs from 12 New World alphavirus family members. Only the peptides containing the tandem motifs are shown.

(G) Representative images of single or combined effect of USP10 peptide and VEEV nsP3 peptide on EEEV and WEEV nsP3 condensate formation. Scale bar, 10 μm.

(H) Dual luciferase assay to analyze reporter activity with the indicated alphaviral nsP3s in HEK293T cells. Error bar indicates ± SEM with n=3 biological replicates.

(I) Dual luciferase assay to analyze reporter activity with MCP control and MCP-VEEV nsP3 in FXR triple KO U2OS cells. Error bar indicates ± SEM with n=3 biological replicates.

(J) A working model depicting the tug of war of the nsP3 condensates and SGs, and the general anti-alphavirus strategy by disrupting the nsP3-G3BP/FXR co-condensates. Semliki Forest virus (SFV) nsP3 co-condenses with the host SG key node protein G3BP, which is critically essential for the intracellular propagation of SFV. Upon SFV infection, the incoming viral RNAs in host cells can lead to the assembly of liquid-like stress granules, in which G3BP acts as the central node and coordinates with other binders (Caprin1, UBAP2L, NUFIP2, etc.) to sequester and shut down the translation of viral RNAs. Furthermore, the SG-sequestered viral RNAs are sensitive to cellular RNase-mediated degradation, which also contributes to SFV restriction (left panel). To overcome the limitation of host SGs, SFV utilizes its nsP3 protein to associate with G3BP, transforming the high-valent SGs into low-valent gel-like nsP3 condensates. The nsP3 condensates support SFV replication by de-repressing viral RNA translation and protecting viral RNA from host RNase (middle panel). Given the pivotal role of SFV nsP3-G3BP co-condensation, targeting SFV nsP3-G3BP interaction via G3BP-competing valence-capping peptides can disrupt nsP3 condensates and inhibit SFV replication. The concept of inhibiting the G3BP-dependent alphaviruses via disrupting nsP3-G3BP co-condensates can be generalizable to the FXR-dependent alphaviruses by targeting nsP3-FXR co-condensation. Thus, nsP3 condensation-targeted therapy can be a promising intervention against alphavirus infection (right panel). Statistical significance in (H) and (I) was assessed using an unpaired, two-tailed *t* test.

## Experimental model and study participant details

### Cell culture

Human osteosarcoma U2OS cells, human embryonic kidney 293T (HEK293T) cells, and baby hamster Syrian kidney (BHK-21) cells were obtained from the National Collection of Authenticated Cell Cultures (Cell Bank, Chinese Academy of Science), cultured in DMEM medium (Invitrogen) supplemented with 10% fetal bovine serum (Cellmax) and 100 U ml^−1^ penicillin and streptomycin. All cell lines were cultured in a humidified incubator at 37 °C with 5% CO_2_. Cell cultures were routinely tested for mycoplasma contamination using a MycoBlue Mycoplasma Detector kit (Vazyme).

## Method details

### Plasmid construction

A human codon-optimized SFV nsP3 coding sequence was inserted into the pEGFP-C3 backbone or an in-house modified pCDH-CMV-mCherry vector for mammalian cell expression and an in-house modified pMal-5x-TEV-His backbone for E. coli expression using the ClonExpress II One Step Cloning Kit (Vazyme). GFP-SFV nsP3 mutants, including Macro/AUD/HVD-deletion and AUD only, C262A/C264A (2CA), R272/275/277A (3RA), R272/275/277E (3RE), F451A (1FA), F468A (1FA), F451A/F468A (2FA) were constructed by PCR with Phanta Max Super-Fidelity DNA Polymerase (Vazyme). To create SFV polyprotein and replicon constructs, the sequences of SFV polyproteins including nsP23, nsP123, nsP1234 and nsP123(3RE)4 were PCR amplified and mutated from the pCMV-SFV4-INT (EGFP-nsP3) construct and inserted into an EGFP-deleted pEGFP-C1 backbone. For *E.coli* expression of gfp-SFV nsP3 FUS_LCD, the FUS_LCD sequence was PCR amplified from HEK293T cDNA and inserted into the SFV nsP3 coding region to substitute the AUD domain in pMal-5x-TEV-His vector. The SFV nsP3_FUS_LCD (G156E) construct was mutated via PCR using the WT construct as template. *E. coli* codon-optimized EEEV/WEEV/VEEV nsP3 coding sequences were synthesized by Tsingke and inserted into pEGFP-C1 for mammalian expression and the pMal-5x-TEV-His backbone for *E. coli* expression. For lentiviral nsP3 constructs, CMV-GFP-SFV/EEEV/WEEV/VEEV nsP3 coding sequences were assembled with a PspXⅠ /EcoR Ⅰ -double digested phND2-N174 construct (Addgene). Coding sequences of G3BP1, Caprin1, USP10, USP10 (F10A), ATXN2, UBAP2L, NUFIP2, YBX1, RNase A/3/5/L, EXOSC10, XRN1, Onconase, and SAMHD1 were PCR-amplified either from the HEK293T cDNA or from a human whole genome ORF library (Thermo Fisher) and assembled with the pEGFP-C1 vector for GFP-fusion expression or the pmCherry-C1 vector for mCherry-fusion expression in cells. Small peptide fragments of Caprin1/USP10/UBAP2L/NUFIP2/SFV nsP3/VEEV nsP3 were amplified from the corresponding full-length proteins and inserted into the pCDH-CMV-mCherry vector. G3BP1 ΔNTF2L/ΔRBD mutants were mutated by PCR from pEGFP-C1-G3BP1 WT construct. *E. coli* codon-optimized G3BP1 and Caprin1 were synthesized by Tsingke and General Biol, respectively, and were inserted into the in-house modified pGEX-6p-1-TEV vector for *E. coli* expression. GST-2×FGDF plasmid was assembled by inserting 2×FGDF into the pGEX-6p-1-TEV vector. The MCP coding sequence was inserted into the gfp-deleted pEGFP-C1 vector, mCherry or 3×HA tag was further assembled in-frame at the downstream of MCP. The customed vectors were used for MCP-mCherry-G3BP1 and MCP-HA-SFV/EEEV/WEEV/VEEV nsP3 construction. For the firefly luciferase-MS2 reporter construction, the firefly luciferase cassette and 12×MS2 coding sequence were simultaneously assembled with the gfp-deleted pEGFP-C1 backbone using the 2×MultiF Seamless Assembly Mix (Abclonal). TetOn-TurboID vector was modified from the pCDH-CMV vector by replacing the CMV promoter with a TetOn promoter consisting of 7 repeats of tetO elements and one minimal CMV promoter and inserting the TurboID-gfp cassette downstream the promoter. G3BP1 coding sequence was inserted into the backbone to create the TetOn-TurboID-GFP-G3BP1 plasmid. Lenti-APEX2-GFP-SFV nsP3 plasmid was constructed by assembling the APEX2, GFP, SFV nsP3 coding sequences in turn into the PspXI/EcoRI-double digested phND2-N174 construct (Addgene). To make the lenti-TetOn-SunTag-nsP4-T2A-Puro-12×MS2 construct, 24×SunTag sequences were PCR amplified using pcDNA4TO-24xGCN4_v4-kif18b-24xPP7 as template, which a kind gift from Weirui Ma (Zhejiang University, China). The 24xSunTag coding sequence, along with PCR-amplified nsP4 CDS, T2A-Puro, and 12×MS fragments were assembled sequentially into a lentiviral vector containing a TetOn promoter. For GFP-VEEV nsP3 FXR-binding mutants, inverse PCR was performed using pEGFP-C1-VEEV nsP3 WT as template. All constructs were confirmed by Sanger sequencing.

### Chemical treatment

To induce stress granule assembly, sodium arsenite solution (Merck) was added into the culture medium at 500 μM for 1 hour. To assess the effect of 1,6-Hexanediol (Sigma-Aldrich) on nsP3 foci, gfp-nsP3 stable U2OS cells were grown in a 20 mm glass bottom culture dish (NEST) 24 hours before the treatment. A final concentration of 3% 1,6-Hexanediol was included in the medium and the live-cell images were taken immediately after addition of 1,6-Hexanediol. To observe whether nsP3 foci are associated with cell membrane-derived vesicles, 10 μM Dil (Thermo Fisher) was incubated with cells for 30 min at 37 °C before fixation. For the qPCR analysis of SFV replication and the nsP3 foci disassembly assay with compounds G3Ib and G3Ib’ (kind gifts from Dr. J. Paul Taylor, St. Jude Children’s Hospital, Memphis), HEK293T or U2OS cells were pre-treated with the indicated compounds (50 μM) or DMSO vehicle for 2 hours before SFV infection for qPCR or fixation for immunofluorescence. For SFV infection tests, the compounds were allowed to be maintained in the medium, and the infected cells were further incubated until harvesting.

### Cell transfection

Transfection of constructs into U2OS, HEK293T, and BHK-21cells was performed using Hieff Trans Liposomal Transfection Reagent (Yeasen) according to the manufacturer’s instructions. Briefly, the DNA constructs and transfection reagent were individually diluted in Opti-MEM medium (Invitrogen) and mixed at a DNA (μg): transfection reagent (μl) ratio of 1: 2. After incubation at room temperature for 15 min, the DNA-liposome complex was added into the cell culture. The medium was removed at 4 hours post-transfection, and the cells were allowed to grow for another 24 hours for analysis or 48 hours for lentiviral production.

### Recombinant virus production and infection

The SFV genome encoding cDNA construct (pCMV-SFV4-INT) was a kind gift from Dr. Andres Merits, University of Tartu, Estonia^57^. The SFV genome cDNA was inserted downstream of a CMV promoter to allow mammalian expression, and a rabbit β-globin intron was included in the Capsid-encoding cassette to ensure well-tolerance in *E.coli*. To generate gfp-/APEX2-recombinant SFV cDNA constructs (pCMV-SFV4-INT (EGFP-nsP3)/(APEX2-nsP3)), the gfp and APEX2-encoding sequences were inserted in-frame with the nsP3 coding region and located between nsP3 Leu 405 and Glu 406. For SFV-2CA/3RE, the 2CA and 3RE mutations were introduced into the SFV genomic DNA construct via inverse PCR. To create the recombinant SFV-SunTag-Luciferase, the 5×SunTag arrays, a firefly luciferase CDS and a minimal SFV subgenomic promoter sequence (GTTATACACCTCTACGGCGGTCCTAGATTGGTGCGTTAA) were inserted into the spacer region between the non-structural polyprotein and the structural polyprotein via seamless assembly. To produce WT, gfp-, APEX2-, or Sun Tag-Luciferase-fused infectious SFV virion, the SFV genome cDNA clone was transfected into BHK-21 cells. At 48 hours post-transfection, the culture supernatant was collected and centrifuged at 12,000 rpm for 15 min at 4 °C to remove cell debris. After filtering through a 0.45 μm PVDF filter, the virus-containing medium was aliquot and stored at -80 °C until use. A serial dilution of the virus-containing medium was used for plague assay in BHK-21 cells to determine viral titers. To infect U2OS and HEK293T cells, SFV was included in FBS-free DMEM at a multiplicity of infection (MOI) 0.1 and incubated with cells at 37 °C for 1 hour with gentle agitation every 15 min. After infection, cells were further incubated with fresh pre-warmed complete DMEM until harvest. GFP-Venezuelan equine encephalitis virus (VEEV TC83 strain)^58^ was a kind gift from Dr. Wenchun Fan, Zhejiang University, China. U2OS WT and FXR1/FXR2/FMR1 triple KO cells were infected at a MOI of 0.1.

### Lentiviral transduction and generation of stable cell lines

gfp-SFV/EEEV/WEEV/VEEV nsP3-and mCherry-SFV nsP3-expressing lentiviruses were produced by co-transfecting the gfp-SFV/EEEV/WEEV/VEEV nsP3 or the mCherry-SFV nsP3-encoding transfer plasmids (CMV promoter) with psPAX2 (Addgene) and pMD2.G (Addgene) plasmids into HEK293T cells. For generating scFv-GFP-expressing lentivirus, pHR-scFv-GCN4-sfGFP-GB1 was co-transfected with psPAX2 and pMD2.G plasmids into HEK293T cells. To produce lentiviruses expressing inducible gfp-TurboID-control/G3BP1, the TetOn-gfp-TurboID-control/ G3BP1 constructs were co-transfected with psPAX2 and pMD2.G plasmids into HEK293T cells. A fourth FUW-M2rtTA plasmid (Addgene) expressing the reverse tetracycline-controlled transactivator (rtTA) was also co-transfected with the above-mentioned plasmids. To produce SunTag-nsP4-inducibly expressing lentivirus, the construct lenti-TetOn-5×SunTag-nsP4-T2A-Puro-12×MS2 was co-transfected with psPAX2, pMD2.G and FUW-M2rtTA plasmids into HEK293T cells. The lentivirus-containing medium was collected after 48 hours and passed through a 0.45 μm filter. For generation of gfp-alphaviral nsP3 stable U2OS cell lines and TetOn-gfp-TurboID-Control/G3BP1 inducible U2OS cell lines, U2OS cells were infected with the above-mentioned lentiviruses for 1 hour and continued to grow in a fresh medium for 47 hours. At 48 hours post-infection, the top 10% of GFP^+^ alphaviral nsP3 expressing cells were sorted using a Sony MA900 cytometer. For sorting TetOn-gfp-TurboID-Control/G3BP1 inducible U2OS cell lines, the transduced cells were treated with 1 μg/ml doxycycline (Solarbio) for 24 hours to induce the expression of gfp-TurboID-Control/G3BP1 proteins before being subjected to FACS. For generation of scFv-GFP stable or scFv-GFP/mCh-SFV nsP3 double stable U2OS cell lines, U2OS cells were infected with the scFv-GFP-expressing lentivirus alone or in combination with the mCherry-SFV nsP3-expressing lentivirus. For generation of SunTag-nsP4-MS2 reporter U2OS cell line, U2OS cells were infected with the scFv-GFP-expressing lentivirus and the TetOn-SunTag-nsP4-T2A-Puro-MS2-expressing lentivirus. To enrich the SunTag-nsP4-MS2 reporter-positive cells, the transduced cells were pre-treated with 1 μg/ml doxycycline for 48h and selected with 0.5 μg/ml puromycin for 21 days in the presence of 1 μg/ml doxycycline. The SunTag-nsP4 reporter/mCh-SFV nsP3 double stable U2OS cells were created by transducing the puromycin-selected SunTag-nsP4 reporter U2OS cells with the mCh-SFV nsP3-expressing lentivirus.

### CRISPR-Cas9 knockout cell lines

Guide RNAs targeting human G3BP1, G3BP2, FXR1, FXR2, and FMR1 were either designed as previously described^24^ or custom-designed using the online Benchling CRISPR Guide RNA Design Tool. The guide RNA oligos were annealed and ligated with the pSpCas9(BB)-2A-Puro (PX459) V2.0 (Addgene) backbone pre-digested with BbsI (NEB). To create G3BP1/2 double knockout U2OS/HEK293T cells and FXR1/FXR2/FMR1 triple knockout U2OS cells, WT U2OS or HEK293T cells were plated in a 6-well plate and transfected with the corresponding plasmids simultaneously for 24 hours. Cells were selected by incubating in DMEM containing 1 μg/ml Puromycin (Biofroxx) for 72 hours. To reduce false positive clones, HEK293T cells, of note, were selected in low confluency by re-plating cells into a 10 cm dish. The selection medium was routinely changed every other day. After antibiotic selection, the survived cells were allowed to recover for 24 hours, followed by diluting into 96-well plates at a concentration of two cells per well. The cells were kept growing for 14-21 days and each single cell clone was checked for protein knockout via immunofluorescence or Western blotting.

### Recombinant protein expression and purification

*E. coli* Rosetta 2(DE3) cells were used to express all proteins. Briefly, the bacteria were grown to OD600 of 0.8 at 37 °C and induced by 1 mM Isopropyl β-D-thiogalactoside (IPTG) (JSENB) at 16 °C for 16 hours. Cells were harvested at 8,000×g for 10 min and resuspended in a high salt lysis buffer containing 50 mM HEPES-NaOH, pH 7.5, 600 mM NaCl, 1 mM DTT (Biofroxx) and 1 mM PMSF (Alpha Diagnostic). Cells were lysed on ice via ultrasonication for a total of 15 min with 2 s of power on and 5 s of power off, at 20 % amplitude. Cell lysates were then centrifuged at 18,000×g for 15 min at 4 °C. For purification of G3BP1, Caprin1 and GST-2×FGDF, 15 ml of the post-cleared supernatant was loaded in a packed GST column with 5 ml glutathione resin (Genscript). The resin was extensively washed with the high salt lysis buffer for a total of 10 column volumes. Coomassie blue-G250 was used to examine the flow-through for sufficient washing. The proteins were eluted with 10 ml of lysis buffer containing 10 mM L-Glutathione reduced (Sigma-Aldrich). The eluate was incubated with home-made TEV protease and 1 μg/ml RNase A overnight at 4 °C to remove the GST tag (except for GST-2×FGDF, in which the GST moiety was kept fusion with the 2×FGDF peptide to increase valency) and RNA contamination. For purification of mCherry, mCherry-G3BP1, gfp-SFV nsP3 WT and its mutants, cells were collected and lysed with the high salt lysis buffer containing 20 mM imidazole (Sigma-Aldrich). The proteins were captured with High Affinity Ni-Charged Resin FF (Genscript), and non-specific proteins were washed with the 20 mM imidazole in lysis buffer. After 10 column volume wash, proteins were eluted with 250 mM imidazole in lysis buffer. To remove the N-terminal solubilizing tag of MBP and reduce RNA contamination, home-made TEV protease and 1 μg/ml RNase A were added into the eluate and incubated overnight at 4 °C. The next day, proteins were filtered through 0.45 μm filters and were subjected to size exclusion chromatography using ÄKTA pure (Cytiva) with a HiLoad 16/600 Superdex 200 pg column (Cytiva). The purified proteins were pooled and concentrated using 10 kDa MWCO Amicon Ultra centrifugal filter (Millipore). Protein concentration was determined by reading A280 absorbance on a NanoDrop One spectrophotometer (Thermo Fisher) and calculated according to the Beer-Lambert equation, c=A280/εL, where c is the protein concentration in molarity, ε is the molar extinction coefficient of the protein and L is the path length. Aliquots of proteins were flash-frozen using liquid nitrogen and stored at -80 °C.

### SFV nsP3 antibody generation

The rabbit polyclonal antibody of nsP3 was raised as previously reported^33^. Briefly, the SFV nsP3 coding sequence was PCR-amplified from the pEGFP-C3-SFV nsP3 construct and inserted in frame with the N-terminal 6x His tag of pET-28a. Stop codon was added to block the C-terminal 6x His tag. The N terminal His-SFV nsP3 plasmid was transformed into *E.coli* Rosetta cells and the recombinant His-SFV nsP3 protein was induced by 1 mM IPTG at 16 °C overnight. The cells were resuspended in lysis buffer (50 mM Tris-HCl, pH 7.5, 400 mM NaCl, 1 mM DTT, 1 mM PMSF) and lysed via ultrasonication on ice for 15 min, with 2 s-on and 5 s-off each cycle. The His-tagged nsP3 was then affinity-purified using the High Affinity Ni-Charged Resin FF resin and further purified by ÄKTA pure with a HiLoad 16/600 Superdex 200 pg column. The purity of the recombinant protein was examined via SDS-PAGE. The purified His-SFV nsP3 protein was then used as the immunogen to immunize two New Zealand White rabbits via HUAbio. The antisera of nsP3 were tested by Western blot preliminarily, and the antibody of nsP3 was further subtracted from the antisera by the Protein A/G purification. The utility and specificity of the purified antibody of nsP3 was verified by Western blot.

### Protein fluorescence labeling

Purified proteins were concentrated to 2 mg/ml in HEPES buffer pH 7.5 containing 600 mM NaCl. 0.1 mM iFluor 488 succinimidyl ester (ATT Bioquest) or XFD647 NHS Ester (ATT Bioquest) was incubated with 1 ml protein solution on a rotator at room temperature for 1 hour in the dark. The reaction was quenched with 20 mM Tris-HCl buffer pH 7.5 for 30 min. To remove free fluorescent dye, protein solution was passed through a HiLoad 16/600 Superdex 200 pg column on ÄKTA pure and the dye-labeled proteins were collected and concentrated. Protein labeling efficiency was measured by reading absorbance at 650 nm using NanoDrop One. For imaging assays, the fluorophore-conjugated proteins and the unlabeled proteins were pre-mixed at a molar ratio of 1:20. In the case of imaging G3BP1 protein *in vitro*, mCherry-fused G3BP1 was mixed with untagged G3BP1 at a molar ratio of 1:20, whereas gfp-fused SFV nsP3 protein was imaged directly without mixing with untagged SFV nsP3 protein.

### *In vitro* transcription

DNA fragment containing a 5’-T7 promoter sequence (TAATACGACTCACTATAGGGAG) and nsP3 coding sequence was PCR amplified and purified using the FastPure Gel DNA Extraction Mini Kit (Vazyme). RNA was *in vitro*-transcribed from 500 ng T7-nsP3 linear DNA fragment dissolved in RNase-free water with the T7 High Yield RNA Transcription Kit (Vazyme) and purified with the RNA Clean & Concentrator-5 kit (Zymo). For transcription of Cy5-labeled nsP3 RNAs, Cyanine 5-UTP (Enzo Life Science) was additionally included in the *in vitro*-transcription reaction at a final concentration of 1 mM.

### Cell fractionation

gfp-nsP3 stable U2OS cells and gfp-SFV infected U2OS cells were homogenized in hypotonic buffer (20 mM Tris-HCl pH 7.4, 1.5 mM MgCl2, 10 mM KCl) supplemented with Protease Inhibitor Cocktail (MCE) by 20 passages through a 1-mL syringe, followed by centrifugation at 1,500× g for 5 min at 4 °C to remove nuclei. The supernatant contained the crude cytoplasmic fraction. The crude cytoplasmic fraction was centrifuged at 5,000×g at 4 °C for 10 min. After centrifugation, the supernatant (S5) was carefully transferred to a new tube and incubated with or without 1% Triton X-100 on ice for 10 min. The S5 fraction was further centrifuged at 20,000×g at 4 °C for 15 min. The post-centrifuged supernatant (S20) and pellet (P20) were separated and analyzed via immunoblot.

### Immunoprecipitation

For co-immunoprecipitation assays, cells were pelleted at 1,500×g for 5 min at 4 °C and lysed in ice-cold IP buffer (20 mM Tris-HCl, pH 7.4, 150 mM NaCl, 1% Triton X-100 and 10% glycerol) supplemented with 1×EDTA-free Protease Inhibitor Cocktail for 30 min on ice. Cell lysates were cleared by centrifugation at 12,000 rpm for 10 min at 4 °C. 10% of the whole cell lysates were reserved as the input by boiling in the SDS loading buffer at 95 °C for 10 min. To precipitate gfp or mCherry-tagged proteins, 1 μg per sample of home-made GST-gfp-nanobody or GST-mCherry-nanobody was conjugated to glutathione agarose resin by incubating on a rotator at room temperature for 1 hour. The nanobody-conjugated GST beads were mixed with the cell lysates and rotated overnight in a cold room at 4 °C. On the next day, the beads were washed extensively with the IP buffer three times, each time on a rotator for 10 min at room temperature. The precipitants were recovered by resuspending and boiling the beads in 2×SDS loading buffer. For the competitive IP of G3Ib and G3Ib’, HEK293T cells were co-transfected with gfp-G3BP1 and HA-SFV nsP3 for 24 h and lysed in ice-cold IP buffer. The compounds were then mixed into the lysates at the indicated concentrations before further subjecting the lysates to IP using the gfp-Nanobody. For immunoprecipitating biotinylated proteins in the proximity labeling assays, cells were lysed in 1×EDTA-free Protease Inhibitor Cocktail supplementing RIPA buffer (20 mM Tris-HCl, pH 7.4, 150 mM NaCl, 1% Triton X-100, 0.5% sodium deoxycholate, 0.1% sodium dodecyl sulfate (SDS)). For the APEX2 labeling cells, 10 mM sodium ascorbate and 5 mM Trolox were additionally included in the RIPA lysis buffer. The APEX2 labeling cell lysates were further dialyzed against fresh RIPA buffer using 10K MWCO Slide-A-Lyzer G3 dialysis cassettes (Thermo Fisher) to remove excessive free biotin-phenol. Both the TurboID and APEX2-mediated biotinylated proteins were enriched using MyOne Streptavidin C1 Dynabeads (Invitrogen) on a rotator overnight at a cold room at 4 °C. After washing the magnetic beads once with 1 M NaCl and three times with RIPA buffer, proteins were eluted by boiling in 2×SDS loading buffer supplemented with 2 mM biotin at 95 °C for 10 min.

### Immunoblot

For regular immunoblot assays, cells were directly lysed and boiled in 2×SDS loading buffer. Cell lysates and immunoprecipitants from co-IP assays were separated on FuturePAGE 4%-20% gels (ACE). Proteins were transferred onto nitrocellulose membranes (Pall). Membranes were blocked with 5% non-fat milk in Tris-buffered saline with 0.1% Tween-20 (TBST), followed by incubation with primary antibodies at 4 °C in a cold room overnight. Membranes were then washed three times with TBST for 5 min each time on an orbital shaker. Membranes were incubated with IRDye 680RD or IRDye 800CW secondary antibodies (LI-COR) for 1 hour in the dark and further washed three times before visualization using the ChemiDoc MP imaging system (Bio-Rad).

### Immunofluorescence

U2OS cells were grown on coverslips in a 24-well plate and fixed with 4% paraformaldehyde (PFA) (Leagene) in PBS for 15 min at room temperature. After washing with PBS once, cells were permeabilized with 0.5% Triton X-100 (Sigma-Aldrich) in PBS for 15 min. 1% BSA in PBST (PBS+ 0.02% Tween 20 (Sigma-Aldrich)) was used to block nonspecific binding in cells for 30 min at room temperature. Primary antibodies were diluted in PBST containing 1% BSA and incubated with cells in a humidified chamber at 4 °C overnight. Cells were washed three times with PBST and incubated with Alexa Fluor 488/555/647-conjugated secondary antibodies (Thermo Fisher) at room temperature in the dark for 1 hour. After incubation, cells were washed three times again with PBST and coverslips were mounted on slides with DAPI (Thermo Fisher) and stored at 4 °C.

### Single molecule fluorescence in situ hybridization (smFISH)

SmFISH probe sequences were designed using the Stellaris RNA FISH Probe Designer software (LGC Biosearch Technologies) against SFV genomic RNA and Quasar 670-labeled oligonucleotide probes were synthesized by General Biol. For visualization of SFV/SFV-SunTag-Luc gRNA, cells were grown on coverslips in a 24-well plate, infected with SFV/ SFV-SunTag-Luc for 12 hours and fixed with 4% paraformaldehyde (PFA) in PBS for 15 min at room temperature. Fixed cells were washed once with PBS and incubated with ice-cold 70% ethanol for 1 hour at 4 °C, then incubated with Buffer A (10% formamide (Sigma-Aldrich) in nuclease-free 2×SSC) for 5 min and hybridized with a pool of 25 Quasar 670-labeled oligonucleotide probes in hybridization buffer (10% dextran sulfate (Sigma-Aldrich) and 10% formamide in 2×SSC) at 37 °C for 16 hours. The next day, cells were washed twice with Buffer A every 30 min at 37 °C and once with nuclease-free 2×SSC for 5 min. Coverslips were mounted on slides with DAPI (Thermo Fisher) and stored at 4 °C. For FISH probe sequences, see Table S4.

### Sequential immunofluorescence and smFISH

For concurrent visualization of G3BP1 and SFV genomic RNA, cells were infected with GFP-SFV for 12 hours and fixed with 4% PFA for 15 min at room temperature. Cells were permeabilized and blocked with blocking buffer (20 mM Tris-HCl, pH 7.5, 150 mM NaCl, 0.5% (w/v) BSA, 0.5% (v/v) Triton X-100 in nuclease-free water) for 1 hour. Cells were then incubated with primary and secondary antibodies as in normal immunofluorescence assay. After incubation with secondary antibody, cells were washed with PBS and fixed with 4% PFA once more for 10 min at room temperature. Cells were washed once with Buffer A (10% formamide in nuclease-free 2×SSC). SFV genomic RNA probe pools were incubated with cells in hybridization buffer (10% dextran sulfate and 10% formamide in 2×SSC) at 37 °C for 16 hours. Following hybridization, cells were washed twice with Buffer A every 30 min at 37 °C and once with nuclease-free 2×SSC for 5 min. Thereafter, coverslips were mounted on slides with DAPI (Thermo Fisher) and stored at 4 °C. For FISH probe sequences, see Table S4.

### *In vitro* LLPS assay

Protein aliquots were thawed at room temperature and shortly centrifuged for 30 s to pellet aggregates. All *in vitro* LLPS assays were assembled in 0.2 ml microtubes at room temperature unless otherwise noted. Briefly, before phase separation, the *in vitro*-transcribed RNAs were diluted in HEPES-NaOH buffer pH 7.5 without salt. The NaCl concentration of protein solutions were adjusted to 300 mM. To initiate LLPS of nsP3 alone, 1 μl of nsP3 in HEPES buffer with 300 mM salt was mixed with 1 μl salt-free HEPES buffer to make the final concentration of NaCl to be 150 mM. For co-phase separation of nsP3, G3BP1 and *in vitro*-transcribed RNA, 1 μl of nsP3 in HEPES buffer with 300 mM salt was pre-mixed with 1 μl of G3BP1 in the same salt buffer, followed by inclusion of 2 μl of RNA in the salt-free HEPES buffer. For nsP3 and Caprin1 competition LLPS assay, nsP3, Caprin1 and G3BP1 were pre-mixed in HEPES buffer with 300 mM NaCl, and an equal volume of *in vitro*-transcribed RNA in the salt-free HEPES buffer was added at last to initiate LLPS. Immediately after the onset of LLPS, 1 μl of the reaction was loaded on a clean glass slide, flanked by two parallel double-sided adhesive tapes. A cover glass was gently placed on top of the tapes to assemble a slit for visualization of LLPS. The assembled chambers were then inverted to allow droplets to settle on the coverslip by gravity for 1 min. For all DIC microscopy of LLPS, images were captured on an Olympus IX73 wide field fluorescence microscope using a UPLXAPO 60× oil immersion objective lens (NA 1.42) and a Hamamatsu ORCA-Flash4.0 LT3 sCMOS camera. For acquisition of multi-color fluorescent LLPS, images were captured using an Olympus FV3000 inverted laser scanning confocal microscope equipped with a UPLXAPO 60× oil immersion objective (NA 1.42).

### Laser scanning confocal microscopy

All single z-slice images of fixed U2OS cells from immunofluorescence and smFISH were captured using an Olympus FV3000 inverted laser scanning confocal microscope equipped with a UPLXAPO 60× oil immersion objective lens (NA 1.42), with the exception that the SFV-SunTag-Luc infected U2OS cells were captured with an Olympus IXplore SpinSR spinning disk confocal microscope equipped with a 100× UPLAPO100XOHR oil immersion objective (NA 1.5). For the sequential smFISH and IF assay, z-stack images were taken at a software-recommended step size. All images were taken under the sequential line scanning mode to minimize fluorescence bleed-through. For image quantification, acquisition parameters such as laser intensity and PMT voltage were identical for all samples in parallel. The 16-bit raw images were exported using the FV31S-SW software and further processed in ImageJ (Fiji).

### Live-cell imaging

For examining nsP3 foci fusion *in vivo*, gfp-nsP3 stable U2OS cells were grown in a 20 mm glass bottom culture dish for 24 hours. Imaging of cells was performed on a Nikon Eclipse Ti2 Spinning Disk Field Scanning confocal microscope equipped with a 60×Apochromat TIRF oil objectives (NA 1.49), in a cage incubator (Okolab) heated to 37 °C with humidified 5% CO_2_. For imaging 1,6-Hexanediol treated gfp-nsP3 stable U2OS cells, the dish lid was kept open during experiment and 1,6-Hexanediol-containing DMEM was dropwise added into cells without disturbing the dish. For all imaging assays, the Nikon perfect focus system was initiated to avoid axial focus fluctuations. For imaging the SunTag-nsP4 translation, the SunTag-nsP4-MS2 reporter U2OS cells were seeded in a 20 mm glass bottom culture dish for 24 hours. The SunTag-nsP4-MS2 mRNA transcription was initiated with 1 μg/ml doxycycline for 2 hours. For monitoring SFV translation in real time, scFv-GFP or scFv-GFP/mCh-SFV nsP3 stable U2OS cells were grown in a 20 mm glass bottom culture dish for 24 hours before being infected with the recombinant SFV-SunTag-Luc for 4-8 hours. The cells were imaged with an Olympus IXplore SpinSR spinning disk (Yokogawa CSU-SoRa) confocal microscope equipped with a 100× UPLAPO100XOHR oil immersion objective (NA 1.5), in a stage top incubator (Takai Hit) at 37 °C with 5% CO_2_. To capture the entire thickness of U2OS cells, 10-20 z stacks were imaged for each field of view (FOV). During multi-color live-cell imaging, a dual-camera (ORCA-Fusion BT sCMOS, Hamamatsu) setup was used to simultaneously capture the signals of GFP-SunTag and mCh-SFV nsP3. The dish lid was allowed to be removed when adding puromycin (100 μg/ml) into the cells.

### *In vitro* and *in vivo* fluorescent recovery after photobleaching (FRAP)

The FRAP experiments were performed on a Nikon Eclipse Ti2 Spinning Disk Field Scanning confocal microscope equipped with a 60×Apochromat TIRF oil objectives (NA 1.49). Five images were acquired prior to photobleaching. For FRAP of *in vitro* nsP3-G3BP1 condensates, a circular ROI of 3-5 μm in diameter was photobleached with 20%-50% laser power for 20 ms using an OBIS LX SF 405 nm direct diode laser module (Coherent). Cellular FRAP of *in vivo* nsP3 foci was performed on the same instrument except that the cells were imaged in a cage incubator with humidified 5% CO_2_ at 37 °C and the size of ROI was set to 2 μm in diameter. Time-lapse images were acquired over 2 min after bleaching. 10-20 spots were bleached for every experiment. FRAP images were analyzed by ImageJ using the FRAP profiler_v2 plugin (developed and released by Dr. Jeff Hardin, University of Wisconsin-Madison). FRAP results were plotted using GraphPad Prism 10.

### *In vitro* RNA protection and time-lapse imaging

For the *in vitro* RNA protection assay, *in vitro* co-LLPS of G3BP1, nsP3 and RNA was induced in a glass bottom 384-well plate (Cellvis) in a total volume of 20 μl. Image acquisition of droplets was immediately started upon gentle addition of 1 μl of RNase A at a final concentration of 1 μg/ml. Acquisition was performed using a Nikon Eclipse Ti2 Spinning Disk (Yokogawa CSU-W1) Field Scanning confocal microscope equipped with a 60×Apochromat TIRF oil objectives (NA 1.49) and a Prime 95B back illuminated sCMOS camera (Teledyne Photometrics). Images were captured every 1 sec and for a total of 15 min.

### Stochastic Optical Reconstruction Microscopy (STORM)

gfp-nsP3 stable U2OS cells or gfp-SFV infected WT U2OS cells were seeded on coverslips and immunolabelled with a gfp nanobody conjugated to Alexa Fluor 647 (1:200, NanoTag Biotechnologies) and a G3BP1 primary antibody (1:500, BD), followed by a secondary antibody conjugated to CF660C (1:2000, Biotium). Two-color super-resolution imaging was performed using the salvaged fluorescence approach^59^ on a custom-built STORM microscope equipped with an oil objective (100× 1.5 NA, UPLAPO100XOHR, Olympus) and a sCMOS camera (ORCA-Fusion BT, C15440-20UP, Hamamatsu) in the presence of imaging buffer. The imaging buffer was freshly prepared immediately before each use. Catalase and glucose oxidase were diluted in a base buffer (44% glycerol, 50 mM Tris-HCl pH 8.0, 10 mM NaCl, 10% glucose) with the addition of βME (Sigma-Aldrich). The final βME concentration was 143 mM. The average excitation intensity of the 642 nm laser (MPB Communications, 2RU-VFL-2000-642-B1R) was approximately 6.75 kW/cm^2^.

### RNA immunoprecipitation (RIP)

For determining RNA binding capacity of nsP3, gfp empty control, gfp-nsP3 WT and 2FA constructs were transfected into 2×10^6^ HEK293T cells for 24 hours, followed by SFV infection for 12 hours. Cells were then washed with ice-cold PBS twice and a minimal amount of ice-cold PBS was added to cell culture dishes after washing. To covalently cross-link RNA and proteins, cells were placed in a UV (254 nm)-cross linker (V-leader) and irradiated with energy mode at 150 mJ/cm^2^. The irradiated cells were transferred on ice and lysed with RIP buffer (20 mM Tris-HCl, pH 7.4, 500 mM NaCl, 10 mM EDTA, 0.5% Triton X-100, 0.1% NP-40, 1 mM DTT and 10% glycerol) containing Protease Inhibitor Cocktail and Recombinant RNase inhibitor (Takara) in RNase-free tubes. After 30 min incubation on ice, cell lysates were centrifuged at 13,000×g for 10 min at 4 °C. 10 % of the post-cleared lysates were transferred to new RNase-free tubes and 3× the sample volume of TRIzol reagent (Invitrogen) was added into the lysate as Input RNA and stored at -80 °C until RNA isolation. For immunoprecipitation, gfp-Trap Magnetic Agarose beads (Chromotek) were incubated with the lysates at 4 °C for 4 hours. After incubation, the magnetic beads were washed with 1 ml high salt wash buffer (20 mM Tris-HCl, pH 7.4, 500 mM NaCl, 10 mM EDTA, 1 mM DTT and 10% glycerol) containing Protease Inhibitor Cocktail and Recombinant RNase inhibitor for 5 min at 4 °C with a total of 5 times. The washed beads were then resuspended in TBS with Recombinant RNase inhibitor and treated with 50 μg/ml Proteinase K (Promega) at 42 °C for 15 min. The supernatant was collected and mixed with 3×volume of TRIzol reagent as IP RNA and stored at -80 °C.

### Fluorescence Polarization assay

For testing RNA binding capacity of SFV nsP3, untagged SFV nsP3 protein stock was two-fold diluted into serial concentrations from 0-8,000 nM in 150 mM HEPES buffer, pH 7.5, and mixed with 20 nM of the FAM-labeled RNA probe in a black 384-well cell culture plates with clear bottom (Beyotime) for 15 min at room temperature in the dark. For assessing RNA binding of nsP3-G3BP1, G3BP1 stock was two-fold diluted into serial concentrations from 0-8,000 nM and mixed with 500 nM SFV nsP3, 20 nM FAM-RNA probe for 15 min at room temperature in the dark. Each value was collected in triplicate. The polarization measurements were performed on a BioTek Synergy Neo2 microplate reader and polarization was calculated with BioTek Gen5 software. The binding curves were fitted to a nonlinear regression one site specific binding model using GraphPad Prism 10.

### MCP-MS2 tethering assay and dual luciferase activity measurement

0.5×10^5^ HEK293T WT or G3BP1/2 dKO cells were plated in a 48-well plate in triplicate and co-transfected with 150 ng MCP-fusion constructs, 50 ng Firefly luciferase-12×MS2 reporter and 2 ng pRL-TK (Promega). For tethering assay in Extended Fig 11b, U2OS FXR tKO cells were used. At 18 hours post-transfection, intracellular luciferase luminescence was measured with a Dual Luciferase Reporter Gene Assay Kit (Yeasen) on a Varioskan LUX microplate reader (Thermo Fisher). Firefly luciferase translation was normalized to the control Renilla luciferase expression.

### RNA isolation and RT-qPCR

Cellular total RNA was isolated using TRIzol reagent according to manufacturer’s instructions. RNA was reverse transcribed using the HiScript III 1st Strand cDNA Synthesis Kit (Vazyme). cDNA was diluted 500-fold with ddH2O and used as templates for qPCR using the ChamQ Universal SYBR qPCR Master Mix (Vazyme) on Bio-Rad CFX Connect Real-Time PCR Detection System. Quantification for all targets was normalized to the control gene β-actin using the 2^^(-ΔΔCt)^ method. For primer sequences, see Table S5.

### Puromycin incorporation assay

For examining cellular bulk translation, cells transfected with mCherry-SG component encoding plasmids with or without gfp-nsP3 plasmid were treated with 10 μg/ml Puromycin for 15 min at 37 °C before being fixed with 4% PFA. Cells were further permeabilized, blocked, and incubated with an anti-Puromycin antibody (Sigma-Aldrich) for immunofluorescence assay.

### TurboID and APEX2 proximity labeling

For the TurboID-mediated proximity labeling, gfp-TurboID-G3BP1 inducible U2OS cells were pre-treated with 1 μg/ml doxycycline for 24 hours, followed by mock or WT SFV infection for 12 hours. 0.2 μg/ml doxycycline was used to induce the expression of gfp-TurboID-Control to a similar level. The culture medium was then replaced with fresh medium containing 50 μM biotin (Sigma-Aldrich) for 10 min to initiate TurboID labeling. Labeling reaction was stopped by moving the cells onto ice and quickly washing the cells five times with ice-cold PBS. For the APEX2-mediated proximity labeling, U2OS cells were infected with recombinant APEX2-SFV for 12 hours, followed by a 30 min incubation of fresh medium containing 500 μM biotin-phenol (Sigma-Aldrich) at 37 °C. To initiate biotinylating reaction, cells were treated with 1 mM H2O2 (Sigma-Aldrich) at room temperature for 1 min. Immediately after labeling, cells were washed three times with freshly prepared quencher solution (10 mM sodium ascorbate (Sigma-Aldrich) and 5 mM Trolox (Sigma-Aldrich) in PBS) to stop labeling.

### Transmission electron microscopy of APEX2-SFV nsP3

Cells were grown on a 20 mm glass bottom culture dish and infected with APEX2-SFV at a MOI of 10. At 12 hours post infection, cells were fixed with 2.5% glutaraldehyde in 0.1 M sodium cacodylate buffer overnight at 4 °C in the dark. Cells were then washed three times with 0.1 M sodium cacodylate buffer and quenched with 20 mM glycine in 0.1 M sodium cacodylate buffer for 5 min on ice. To initiate DAB staining, cells were incubated with freshly prepared 0.5 mg/ml DAB (Sigma-Aldrich) and 10 mM H2O2 in 0.1 M sodium cacodylate buffer for 25-30 min on ice. The DAB solution was then gently removed, and cells were washed five times with 0.1 M sodium cacodylate buffer. Cells were stained with 2% osmium tetroxide and uranyl acetate for 1 hour on ice. After osmium tetroxide staining, cells were dehydrated by sequentially submerging into 20, 50, 70, 90, and 100% (vol/vol) ethanol, and embedded in epoxy resin. Then ultrathin sectioning was carried out using a Leica EM UC7 ultramicrotome. Images were taken with a Talos L120C transmission electron microscope (120 kV) and a bottom-mounted 4k×4k Ceta CMOS camera (Thermo Fisher).

### Mass spectrometry

For identification of G3BP1 and nsP3 interactome using TurboID/APEX2-labeling strategy, cell samples in duplicate were immunoprecipitated with Streptavidin-conjugated magnetic beads and run on a FuturePAGE 4%-20% polyacrylamide gel at 160 V for 10 min. The samples were sliced with a clean single side security blade and were subjected to overnight in-gel digestion with trypsin following reduction and alkylation with DTT and iodoacetamide in 50 mM ammonium bicarbonate at 37°C. For LC-MS/MS analysis, the peptides were separated by a 65 min gradient elution at a flow rate of 0.3 μL/min with the Thermo EASY-nLC1200 integrated nano-HPLC system which is directly interfaced with the Thermo Orbitrap Exploris 480 mass spectrometer. The analytical column was a homemade fused silica capillary column (75 μm ID, 150 mm length; Upchurch, Oak Harbor, WA) packed with C-18 resin (300 A, 3 μm, Varian, Lexington, MA). Mobile phase A consisted of 0.1% formic acid, and mobile phase B consisted of 100% acetonitrile and 0.1% formic acid. The mass spectrometer was operated in the data-dependent acquisition mode using the Xcalibur 4.1 software and there is a single full-scan mass spectrum in the Orbitrap (400–1800 m/z, 60,000 resolution) followed by 20 data-dependent MS/MS scans at 30% normalized collision energy. Each mass spectrum was analyzed using the Thermo Xcalibur Qual Browser and Proteome Discovery for database searching.

### Proteomic data analysis

For label-free quantification analysis of TurboID-G3BP1 and APEX2-nsP3 proteome, raw data of each group were searched using MaxQuant software (v.2.4.9.0). The FASTA file containing the complete human proteome (downloaded from UniProt) as well as commonly observed contaminants was used as the reference for protein identification. The following search parameters were used: digestion was specified as trypsin with up to 2 missed cleavages. Fixed modification was carbamidomethyl[C] and variable modifications were oxidation of methionine and acetyl on protein N termini. Spectra were searched with a mass accuracy of 4.5 ppm for precursors and 20 ppm for fragment ions. The false discovery rate (FDR) was set to 0.01, both at protein and peptide levels, using a reverse database as a decoy. Label-free quantification parameters were set as default. For differential expression analysis, the processed files were analyzed using an online LFQ-Analyst tool^60^ according to the detailed user manual. Briefly, the MaxQuant proteinGroups.txt files were uploaded, adjusted p-value cutoff was set to 0.05 and Log2 fold change cutoff was set to 0.65. The Benjamini Hochberg (BH) method was used for false discovery rate (FDR) correction. Quantified results were visualized using the Volcano tool in Hiplot Pro, a comprehensive web service for biomedical data analysis and visualization.

### Gene Ontology

Gene functional annotation and clustering of the hits acquired from the mass spectrometry were analyzed and profiled using the GO/KEGG cluster profiler tool in Hiplot Pro with a Benjamini-Hochberg multiple testing adjustment under an FDR cutoff of 0.05.

### Molecular docking

Protein_Jprotein docking is used to predict interactions between NTF2L domain of G3BP1 and NUFIP2. The initial structure of the NTF2L/NUFIP2 complex was predicted using AlphaFold3^61^. Rigid docking of NTF2L and NUFIP2 was performed using the ZDOCK 3.0.2^62^. The docking results were visualized and further analyzed using PyMol. Crystal structures of NTF2L bound to peptides 6TA7 (Caprin1), 8TH6 (USP10), and 5FW5 (nsP3), were aligned to the best docking structure of NTF2L and NUFIP2 using the “align” method.

### Word frequency analysis of GO terms (Word Cloud)

A spreadsheet of differential genes identified in TurboID/APEX2-labeling proteomes was loaded in WocEA (v.1.0) software^63^ and run for GO enrichment. The top GO terms and their corresponding enrichment ratio (E-ratio) (denoted by color) and p-value (denoted by font size) were used to create the word cloud layout.

### Estimation of intracellular protein concentration

For absolute intracellular G3BP1 and nsP3 concentration estimation, 1×106 HEK293T cells were infected with gfp-SFV (MOI 0.1) for the indicated time points and directly lysed in 50 μl of 2×SDS loading buffer. To generate a standard calibration curve for directly converting the measured densitometry into absolute protein concentration, recombinant G3BP1 and gfp-SFV nsP3 proteins were two-fold diluted in concentrations from 0.25 μM to 32 μM in 2×SDS loading buffer. 5 μl of each protein standard and cell lysates were run on a FuturePAGE 4%-20% polyacrylamide gel and immunoblotted with either G3BP1 or gfp antibody. ImageJ was used to analyze the band intensity of G3BP1 and gfp-nsP3. The average cell volume of HEK293T was estimated to be 1×10^-12^ L^56^. The absolute concentration of proteins could be calculated as following:

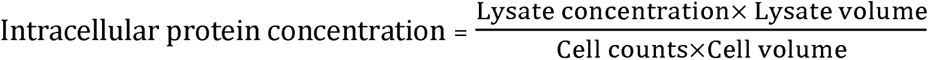

### LLPS threshold concentration measurement *in vivo*

U2OS cells were transiently transfected with various doses of gfp-SFV nsP3 WT and 2FA mutant to achieve a diversified concentration range. All the cells were imaged at the z-slice with the highest fluorescence intensity by adjusting the focus. For all images, identical capture parameters (including laser power, PMT gain, and scanning resolution) were used. All measurements in each experiment were performed on the same confocal microscope. Recombinant gfp-SFV nsP3 protein stock was two-fold diluted to make a serial of eight standards that were used to build a calibration curve. The standard curve was used to convert the absolute fluorescence intensity to absolute molar concentration. To calculate the nsP3 absolute concentration, a circular ROI of 2 μm in diameter with average intensity was drawn in the diffusive gfp area within every single cell expressing gfp-nsP3 WT/2FA using ImageJ. A total of 100 or more cells were measured. The fluorescence intensity of each cell was converted to absolute concentration using the standard curve. For determining the nsP3 LLPS threshold concentration, 100 cells were sorted by their concentrations from high to low and were binned to 10 groups, with each bin containing 10 cells. The bin with more than 5 nsP3 foci-positive cells (defined as cells having at least 3 nsP3 foci) was used to determine the threshold concentration. The mean concentration of the bin was calculated as the threshold concentration, and the bin size was used to calculate the error.

### Colocalization analysis of G3BP1 and SFV genomic RNA

The image stack of G3BP1 and SFV genomic RNA was analyzed in ImageJ using the Z projection Max intensity tool, and colocalization was demonstrated using line profiling.

### Analysis of SunTag translation with puromycin

Time-lapse images of SunTag translation after puromycin treatment were analyzed as maximum intensity projections using Image J. Bleach correction using the histogram matching method in Image J was performed to adjust the intensity attenuation caused by long-term imaging.

### Track analysis of nsP3 foci and SunTag translation

Single particle tracking of nsP3 and SunTag foci was performed on the maximum intensity projection of z-stack images using the TrackMate plugin of Image J. Briefly, foci of translation and nsP3 were first identified by adjusting intensity threshold. The thresholding image was then transformed into a binary image that was used for TrackMate analysis. The Laplacian of Gaussian (LoG) detector was used to detect particles. A simple linear assignment problem (LAP) tracker was used to track the foci trajectory. The results of x, y and t were plotted using Origin2024.

### Stress granule-nsP3 granule competition assay *in vitro*

For analyzing nsP3 and SG formation *in vitro*, 5 μM mCherry-G3BP1 protein was mixed with the indicated concentration of gfp-nsP3 WT/mutants and XFD647-labeled Caprin1 proteins. Phase separation was induced with 100 ng/μl *in vitro*-transcribed nsP3 RNA. Images of the multi-color droplets were taken by an Olympus FV3000 confocal microscope, and the fluorescence intensity of Caprin1 inside nsP3-G3BP1 condensates was measured by ImageJ. Caprin1 intensities were plotted against the nsP3 concentrations, and the fitted curves were generated using GraphPad Prism 10. All assays in parallel were conducted using identical parameters on the same confocal microscope.

### Analysis of stress granule disassembly in nsP3-expressing U2OS cells

To measure stress granule dynamics in the presence of nsP3, U2OS cells were transfected with gfp-SFV nsP3 WT and FA mutants for 24 hours and treated with SA for 1 hour to induce stress granule formation. After 1 hour, cells were washed three times with pre-warmed culture medium and further incubated for the indicated time points. Cells were fixed and subjected to immunofluorescent staining using G3BP1 and Caprin1 antibodies. Images of transfected cells at each time point were captured by an Olympus FV3000 confocal microscope and analyzed using ImageJ. The number of cells visible with stress granules (defined as cells that harbored double positive staining foci for G3BP1 and Caprin1) was manually counted in each field of view (FOV). The ratio of SG-positive cells over nsP3-positive cells was calculated, plotted, and fitted using GraphPad Prism 10.

### Line profiling

Line analysis was performed on ImageJ. Briefly, a line was drawn and saved as an ROI in the ROI manager. Images of each channel were stacked, and the line ROI was added to the stack file. Line intensity of each channel was measured using the Plot Profile tool, exported into a CSV file and further plotted using GraphPad Prism 10.

### Partition coefficient measurement

G3BP1 partition *in vitro* was measured using ImageJ. A circular ROI was manually created to include a single G3BP1 droplet, and the mean fluorescent intensity of the ROI was measured. To measure diffusive G3BP1 intensity, the frame was converted into 16-bit image, and the threshold values were manually adjusted to include all G3BP1 droplets in the frame. The image was then converted to a binary image and the intensity sum of the droplets in the frame was measured using the “Analyze particles” tool. The total intensity of the frame was measured using the tool “Measure”. The mean intensity of diffusive G3BP1 in the background could be calculated as:

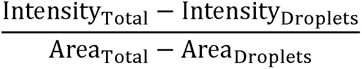

Thus, the partition coefficient of G3BP1 was calculated by taking the ratio of the mean fluorescence intensity inside G3BP1 droplet to the mean fluorescence intensity in the background.

### Sequence alignment and phylogenetic tree of alphaviral nsP3s

Sequences of all alphaviruses nsP3 amino acid sequences were collected and downloaded from UniProt. Multi-sequence alignment was performed in the MEGA 11 software using the ClustalW algorithm^64^. The phylogenetic tree of nsP3s was constructed using the maximum likelihood (ML) method. The alignment result was exported as a FASTA file and visualized using a web service program ESPript 3.0^65^.

### Quantification and statistical analysis

Statistical analyses were performed in GraphPad Prism 10 using either the unpaired Student’s *t* test, one-way ANOVA, or two-way ANOVA test as stated in each figure legend. Quantification data of qPCR assays in bar plots were expressed as means ± SEM, in which each dot represents one value in parallel. Data of FRAP assays in line plots were expressed as means ± SEM. Data of partition coefficient and LLPS threshold measurements in box and whiskers plots were expressed as the first quartile, median, and third quartile for the lower, middle, and upper line, respectively, and minimum to maximum for the down and up extension of whiskers, in which each dot represents one individual cell or droplet *in vitro*. ns, not significant; * *p* < 0.05, ** *p* < 0.01, *** *p* < 0.001 and **** *p* < 0.0001. Data are judged to be statistically significant when *p*< 0.05.

## Reference

1. Sagan, S.M., and Weber, S.C. (2023). Let’s phase it: viruses are master architects of biomolecular condensates. Trends Biochem Sci 48, 229–243. 10.1016/j.tibs.2022.09.008.

2. Borodavka, A., and Acker, J. (2023). Seeing Biomolecular Condensates Through the Lens of Viruses. Annu Rev Virol 10, 163–182. 10.1146/annurev-virology-111821-103226.

3. Banani, S.F., Lee, H.O., Hyman, A.A., and Rosen, M.K. (2017). Biomolecular condensates: organizers of cellular biochemistry. Nat Rev Mol Cell Biol 18, 285–298. 10.1038/nrm.2017.7.

4. Lyon, A.S., Peeples, W.B., and Rosen, M.K. (2021). A framework for understanding the functions of biomolecular condensates across scales. Nat Rev Mol Cell Biol 22, 215–235. 10.1038/s41580-020-00303-z.

5. Geiger, F., Acker, J., Papa, G., Wang, X., Arter, W.E., Saar, K.L., Erkamp, N.A., Qi, R., Bravo, J.P., Strauss, S., et al. (2021). Liquid-liquid phase separation underpins the formation of replication factories in rotaviruses. EMBO J 40, e107711. 10.15252/embj.2021107711.

6. Miyake, T., Farley, C.M., Neubauer, B.E., Beddow, T.P., Hoenen, T., and Engel, D.A. (2020). Ebola Virus Inclusion Body Formation and RNA Synthesis Are Controlled by a Novel Domain of Nucleoprotein Interacting with VP35. J Virol 94. 10.1128/JVI.02100-19.

7. Guseva, S., Milles, S., Jensen, M.R., Salvi, N., Kleman, J.P., Maurin, D., Ruigrok, R.W.H., and Blackledge, M. (2020). Measles virus nucleo- and phosphoproteins form liquid-like phase-separated compartments that promote nucleocapsid assembly. Sci Adv 6, eaaz7095. 10.1126/sciadv.aaz7095.

8. Zhou, Y., Su, J.M., Samuel, C.E., and Ma, D. (2019). Measles Virus Forms Inclusion Bodies with Properties of Liquid Organelles. J Virol 93. 10.1128/JVI.00948-19.

9. Alenquer, M., Vale-Costa, S., Etibor, T.A., Ferreira, F., Sousa, A.L., and Amorim, M.J. (2019). Influenza A virus ribonucleoproteins form liquid organelles at endoplasmic reticulum exit sites. Nat Commun 10, 1629. 10.1038/s41467-019-09549-4.

10. Rincheval, V., Lelek, M., Gault, E., Bouillier, C., Sitterlin, D., Blouquit-Laye, S., Galloux, M., Zimmer, C., Eleouet, J.F., and Rameix-Welti, M.A. (2017). Functional organization of cytoplasmic inclusion bodies in cells infected by respiratory syncytial virus. Nat Commun 8, 563. 10.1038/s41467-017-00655-9.

11. Iserman, C., Roden, C.A., Boerneke, M.A., Sealfon, R.S.G., McLaughlin, G.A., Jungreis, I., Fritch, E.J., Hou, Y.J., Ekena, J., Weidmann, C.A., et al. (2020). Genomic RNA Elements Drive Phase Separation of the SARS-CoV-2 Nucleocapsid. Mol Cell 80, 1078–1091 e1076. 10.1016/j.molcel.2020.11.041.

12. Heinrich, B.S., Maliga, Z., Stein, D.A., Hyman, A.A., and Whelan, S.P.J. (2018). Phase Transitions Drive the Formation of Vesicular Stomatitis Virus Replication Compartments. mBio 9. 10.1128/mBio.02290-17.

13. Risso-Ballester, J., Galloux, M., Cao, J., Le Goffic, R., Hontonnou, F., Jobart-Malfait, A., Desquesnes, A., Sake, S.M., Haid, S., Du, M., et al. (2021). A condensate-hardening drug blocks RSV replication in vivo. Nature 595, 596–599. 10.1038/s41586-021-03703-z.

14. Charman, M., Grams, N., Kumar, N., Halko, E., Dybas, J.M., Abbott, A., Lum, K.K., Blumenthal, D., Tsopurashvili, E., and Weitzman, M.D. (2023). A viral biomolecular condensate coordinates assembly of progeny particles. Nature 616, 332–338. 10.1038/s41586-023-05887-y.

15. De Caluwe, L., Arien, K.K., and Bartholomeeusen, K. (2021). Host Factors and Pathways Involved in the Entry of Mosquito-Borne Alphaviruses. Trends Microbiol 29, 634–647. 10.1016/j.tim.2020.10.011.

16. Kim, A.S., and Diamond, M.S. (2023). A molecular understanding of alphavirus entry and antibody protection. Nat Rev Microbiol 21, 396–407. 10.1038/s41579-022-00825-7.

17. Gotte, B., Liu, L., and McInerney, G.M. (2018). The Enigmatic Alphavirus Non-Structural Protein 3 (nsP3) Revealing Its Secrets at Last. Viruses 10. 10.3390/v10030105.

18. Foy, N.J., Akhrymuk, M., Akhrymuk, I., Atasheva, S., Bopda-Waffo, A., Frolov, I., and Frolova, E.I. (2013). Hypervariable domains of nsP3 proteins of New World and Old World alphaviruses mediate formation of distinct, virus-specific protein complexes. J Virol 87, 1997–2010. 10.1128/JVI.02853-12.

19. Gorchakov, R., Garmashova, N., Frolova, E., and Frolov, I. (2008). Different types of nsP3-containing protein complexes in Sindbis virus-infected cells. J Virol 82, 10088–10101. 10.1128/JVI.01011-08.

20. Lark, T., Keck, F., and Narayanan, A. (2017). Interactions of Alphavirus nsP3 Protein with Host Proteins. Front Microbiol 8, 2652. 10.3389/fmicb.2017.02652.

21. Cristea, I.M., Carroll, J.W., Rout, M.P., Rice, C.M., Chait, B.T., and MacDonald, M.R. (2006). Tracking and elucidating alphavirus-host protein interactions. J Biol Chem 281, 30269–30278. 10.1074/jbc.M603980200.

22. Kril, V., Hons, M., Amadori, C., Zimberger, C., Couture, L., Bouery, Y., Burlaud-Gaillard, J., Karpov, A., Ptchelkine, D., Thienel, A.L., et al. (2024). Alphavirus nsP3 organizes into tubular scaffolds essential for infection and the cytoplasmic granule architecture. Nat Commun 15, 8106. 10.1038/s41467-024-51952-z.

23. Clark, L.E., Clark, S.A., Lin, C., Liu, J., Coscia, A., Nabel, K.G., Yang, P., Neel, D.V., Lee, H., Brusic, V., et al. (2022). VLDLR and ApoER2 are receptors for multiple alphaviruses. Nature 602, 475–480. 10.1038/s41586-021-04326-0.

24. Yang, P., Mathieu, C., Kolaitis, R.M., Zhang, P., Messing, J., Yurtsever, U., Yang, Z., Wu, J., Li, Y., Pan, Q., et al. (2020). G3BP1 Is a Tunable Switch that Triggers Phase Separation to Assemble Stress Granules. Cell 181, 325–345 e328. 10.1016/j.cell.2020.03.046.

25. Sanders, D.W., Kedersha, N., Lee, D.S.W., Strom, A.R., Drake, V., Riback, J.A., Bracha, D., Eeftens, J.M., Iwanicki, A., Wang, A., et al. (2020). Competing Protein-RNA Interaction Networks Control Multiphase Intracellular Organization. Cell 181, 306–324 e328. 10.1016/j.cell.2020.03.050.

26. Guillen-Boixet, J., Kopach, A., Holehouse, A.S., Wittmann, S., Jahnel, M., Schlussler, R., Kim, K., Trussina, I., Wang, J., Mateju, D., et al. (2020). RNA-Induced Conformational Switching and Clustering of G3BP Drive Stress Granule Assembly by Condensation. Cell 181, 346–361 e317. 10.1016/j.cell.2020.03.049.

27. Jain, S., Wheeler, J.R., Walters, R.W., Agrawal, A., Barsic, A., and Parker, R. (2016). ATPase-Modulated Stress Granules Contain a Diverse Proteome and Substructure. Cell 164, 487–498. 10.1016/j.cell.2015.12.038.

28. Panas, M.D., Schulte, T., Thaa, B., Sandalova, T., Kedersha, N., Achour, A., and McInerney, G.M. (2015). Viral and cellular proteins containing FGDF motifs bind G3BP to block stress granule formation. PLoS Pathog 11, e1004659. 10.1371/journal.ppat.1004659.

29. Cho, N.H., Cheveralls, K.C., Brunner, A.D., Kim, K., Michaelis, A.C., Raghavan, P., Kobayashi, H., Savy, L., Li, J.Y., Canaj, H., et al. (2022). OpenCell: Endogenous tagging for the cartography of human cellular organization. Science 375, eabi6983. 10.1126/science.abi6983.

30. Patel, A., Lee, H.O., Jawerth, L., Maharana, S., Jahnel, M., Hein, M.Y., Stoynov, S., Mahamid, J., Saha, S., Franzmann, T.M., et al. (2015). A Liquid-to-Solid Phase Transition of the ALS Protein FUS Accelerated by Disease Mutation. Cell 162, 1066–1077. 10.1016/j.cell.2015.07.047.

31. McCormick, C., and Khaperskyy, D.A. (2017). Translation inhibition and stress granules in the antiviral immune response. Nat Rev Immunol 17, 647–660. 10.1038/nri.2017.63.

32. Burke, J.M., Ratnayake, O.C., Watkins, J.M., Perera, R., and Parker, R. (2024). G3BP1-dependent condensation of translationally inactive viral RNAs antagonizes infection. Sci Adv 10, eadk8152. 10.1126/sciadv.adk8152.

33. Yao, Z., Liu, Y., Chen, Q., Chen, X., Zhu, Z., Song, S., Ma, X., and Yang, P. (2024). The divergent effects of G3BP orthologs on human stress granule assembly imply a centric role for the core protein interaction network. Cell Rep 43, 114617. 10.1016/j.celrep.2024.114617.

34. Case, L.B., Zhang, X., Ditlev, J.A., and Rosen, M.K. (2019). Stoichiometry controls activity of phase-separated clusters of actin signaling proteins. Science 363, 1093–1097. 10.1126/science.aau6313.

35. Banani, S.F., Rice, A.M., Peeples, W.B., Lin, Y., Jain, S., Parker, R., and Rosen, M.K. (2016). Compositional Control of Phase-Separated Cellular Bodies. Cell 166, 651–663. 10.1016/j.cell.2016.06.010.

36. Palchevska, O., Dominguez, F., Frolova, E.I., and Frolov, I. (2023). Alphavirus-induced transcriptional and translational shutoffs play major roles in blocking the formation of stress granules. bioRxiv. 10.1101/2023.07.05.547824.

37. Jayabalan, A.K., Adivarahan, S., Koppula, A., Abraham, R., Batish, M., Zenklusen, D., Griffin, D.E., and Leung, A.K.L. (2021). Stress granule formation, disassembly, and composition are regulated by alphavirus ADP-ribosylhydrolase activity. Proc Natl Acad Sci U S A 118. 10.1073/pnas.2021719118.

38. Panas, M.D., Varjak, M., Lulla, A., Eng, K.E., Merits, A., Karlsson Hedestam, G.B., and McInerney, G.M. (2012). Sequestration of G3BP coupled with efficient translation inhibits stress granules in Semliki Forest virus infection. Mol Biol Cell 23, 4701–4712. 10.1091/mbc.E12-08-0619.

39. Schulte, T., Liu, L., Panas, M.D., Thaa, B., Dickson, N., Gotte, B., Achour, A., and McInerney, G.M. (2016). Combined structural, biochemical and cellular evidence demonstrates that both FGDF motifs in alphavirus nsP3 are required for efficient replication. Open Biol 6. 10.1098/rsob.160078.

40. Schulte, T., Panas, M.D., Han, X., Williams, L., Kedersha, N., Fleck, J.S., Tan, T.J.C., Dopico, X.C., Olsson, A., Morro, A.M., et al. (2023). Caprin-1 binding to the critical stress granule protein G3BP1 is influenced by pH. Open Biol 13, 220369. 10.1098/rsob.220369.

41. Mutso, M., Morro, A.M., Smedberg, C., Kasvandik, S., Aquilimeba, M., Teppor, M., Tarve, L., Lulla, A., Lulla, V., Saul, S., et al. (2018). Mutation of CD2AP and SH3KBP1 Binding Motif in Alphavirus nsP3 Hypervariable Domain Results in Attenuated Virus. Viruses 10. 10.3390/v10050226.

42. Youn, J.Y., Dunham, W.H., Hong, S.J., Knight, J.D.R., Bashkurov, M., Chen, G.I., Bagci, H., Rathod, B., MacLeod, G., Eng, S.W.M., et al. (2018). High-Density Proximity Mapping Reveals the Subcellular Organization of mRNA-Associated Granules and Bodies. Mol Cell 69, 517–532 e511. 10.1016/j.molcel.2017.12.020.

43. Markmiller, S., Soltanieh, S., Server, K.L., Mak, R., Jin, W., Fang, M.Y., Luo, E.C., Krach, F., Yang, D., Sen, A., et al. (2018). Context-Dependent and Disease-Specific Diversity in Protein Interactions within Stress Granules. Cell 172, 590–604 e513. 10.1016/j.cell.2017.12.032.

44. Kedersha, N., Panas, M.D., Achorn, C.A., Lyons, S., Tisdale, S., Hickman, T., Thomas, M., Lieberman, J., McInerney, G.M., Ivanov, P., and Anderson, P. (2016). G3BP-Caprin1-USP10 complexes mediate stress granule condensation and associate with 40S subunits. J Cell Biol 212, 845–860. 10.1083/jcb.201508028.

45. Freibaum, B.D., Messing, J., Nakamura, H., Yurtsever, U., Wu, J., Kim, H.J., Hixon, J., Lemieux, R.M., Duffner, J., Huynh, W., et al. (2024). Identification of small molecule inhibitors of G3BP-driven stress granule formation. J Cell Biol 223. 10.1083/jcb.202308083.

46. Kim, D.Y., Reynaud, J.M., Rasalouskaya, A., Akhrymuk, I., Mobley, J.A., Frolov, I., and Frolova, E.I. (2016). New World and Old World Alphaviruses Have Evolved to Exploit Different Components of Stress Granules, FXR and G3BP Proteins, for Assembly of Viral Replication Complexes. PLoS Pathog 12, e1005810. 10.1371/journal.ppat.1005810.

47. Frolov, I., Kim, D.Y., Akhrymuk, M., Mobley, J.A., and Frolova, E.I. (2017). Hypervariable Domain of Eastern Equine Encephalitis Virus nsP3 Redundantly Utilizes Multiple Cellular Proteins for Replication Complex Assembly. J Virol 91. 10.1128/JVI.00371-17.

48. Laver, J.D., Ly, J., Winn, A.K., Karaiskakis, A., Lin, S., Nie, K., Benic, G., Jaberi-Lashkari, N., Cao, W.X., Khademi, A., et al. (2020). The RNA-Binding Protein Rasputin/G3BP Enhances the Stability and Translation of Its Target mRNAs. Cell Rep 30, 3353–3367 e3357. 10.1016/j.celrep.2020.02.066.

49. Liu, Y., Yao, Z., Lian, G., and Yang, P. (2023). Biomolecular phase separation in stress granule assembly and virus infection. Acta Biochim Biophys Sin (Shanghai) 55, 1099–1118. 10.3724/abbs.2023117.

50. Frolova, E.I., Palchevska, O., Dominguez, F., and Frolov, I. (2023). Alphavirus-induced transcriptional and translational shutoffs play major roles in blocking the formation of stress granules. J Virol 97, e0097923. 10.1128/jvi.00979-23.

51. Foy, N.J., Akhrymuk, M., Shustov, A.V., Frolova, E.I., and Frolov, I. (2013). Hypervariable domain of nonstructural protein nsP3 of Venezuelan equine encephalitis virus determines cell-specific mode of virus replication. J Virol 87, 7569–7584. 10.1128/JVI.00720-13.

52. Hosmillo, M., Lu, J., McAllaster, M.R., Eaglesham, J.B., Wang, X., Emmott, E., Domingues, P., Chaudhry, Y., Fitzmaurice, T.J., Tung, M.K., et al. (2019). Noroviruses subvert the core stress granule component G3BP1 to promote viral VPg-dependent translation. Elife 8. 10.7554/eLife.46681.

53. Yang, Z., Johnson, B.A., Meliopoulos, V.A., Ju, X., Zhang, P., Hughes, M.P., Wu, J., Koreski, K.P., Clary, J.E., Chang, T.C., et al. (2024). Interaction between host G3BP and viral nucleocapsid protein regulates SARS-CoV-2 replication and pathogenicity. Cell Rep 43, 113965. 10.1016/j.celrep.2024.113965.

54. Yi, Z., Pan, T., Wu, X., Song, W., Wang, S., Xu, Y., Rice, C.M., Macdonald, M.R., and Yuan, Z. (2011). Hepatitis C virus co-opts Ras-GTPase-activating protein-binding protein 1 for its genome replication. J Virol 85, 6996–7004. 10.1128/JVI.00013-11.

55. Li, M., Hou, Y., Zhou, Y., Yang, Z., Zhao, H., Jian, T., Yu, Q., Zeng, F., Liu, X., Zhang, Z., and Zhao, Y.G. (2024). LLPS of FXR proteins drives replication organelle clustering for beta-coronaviral proliferation. J Cell Biol 223. 10.1083/jcb.202309140.

56. Milo, R. (2013). What is the total number of protein molecules per cell volume? A call to rethink some published values. Bioessays 35, 1050–1055. 10.1002/bies.201300066.

57. Ulper, L., Sarand, I., Rausalu, K., and Merits, A. (2008). Construction, properties, and potential application of infectious plasmids containing Semliki Forest virus full-length cDNA with an inserted intron. J Virol Methods 148, 265–270. 10.1016/j.jviromet.2007.10.007.

58. Xie, Y., Cao, J., Gan, S., Xu, L., Zhang, D., Qian, S., Xu, F., Ding, Q., Schoggins, J.W., and Fan, W. (2024). TRIM32 inhibits Venezuelan equine encephalitis virus infection by targeting a late step in viral entry. PLoS Pathog 20, e1012312. 10.1371/journal.ppat.1012312.

59. Zhang, Y., Schroeder, L.K., Lessard, M.D., Kidd, P., Chung, J., Song, Y., Benedetti, L., Li, Y., Ries, J., Grimm, J.B., et al. (2020). Nanoscale subcellular architecture revealed by multicolor three-dimensional salvaged fluorescence imaging. Nat Methods 17, 225–231. 10.1038/s41592-019-0676-4.

60. Shah, A.D., Goode, R.J.A., Huang, C., Powell, D.R., and Schittenhelm, R.B. (2020). LFQ-Analyst: An Easy-To-Use Interactive Web Platform To Analyze and Visualize Label-Free Proteomics Data Preprocessed with MaxQuant. J Proteome Res 19, 204–211. 10.1021/acs.jproteome.9b00496.

61. Abramson, J., Adler, J., Dunger, J., Evans, R., Green, T., Pritzel, A., Ronneberger, O., Willmore, L., Ballard, A.J., Bambrick, J., et al. (2024). Accurate structure prediction of biomolecular interactions with AlphaFold 3. Nature 630, 493–500. 10.1038/s41586-024-07487-w.

62. Pierce, B.G., Hourai, Y., and Weng, Z. (2011). Accelerating protein docking in ZDOCK using an advanced 3D convolution library. PLoS One 6, e24657. 10.1371/journal.pone.0024657.

63. Ning, W., Lin, S., Zhou, J., Guo, Y., Zhang, Y., Peng, D., Deng, W., and Xue, Y. (2018). WocEA: The visualization of functional enrichment results in word clouds. J Genet Genomics 45, 415–417. 10.1016/j.jgg.2018.02.008.

64. Tamura, K., Stecher, G., and Kumar, S. (2021). MEGA11: Molecular Evolutionary Genetics Analysis Version 11. Mol Biol Evol 38, 3022–3027. 10.1093/molbev/msab120.

65. Robert, X., and Gouet, P. (2014). Deciphering key features in protein structures with the new ENDscript server. Nucleic Acids Res 42, W320–324. 10.1093/nar/gku316.

