## Supplementary figures and images for "Alphaviral nonstructural protein-host RBP co-condensation as a mechanism to sustain virus replication"

### Supplemental Figures

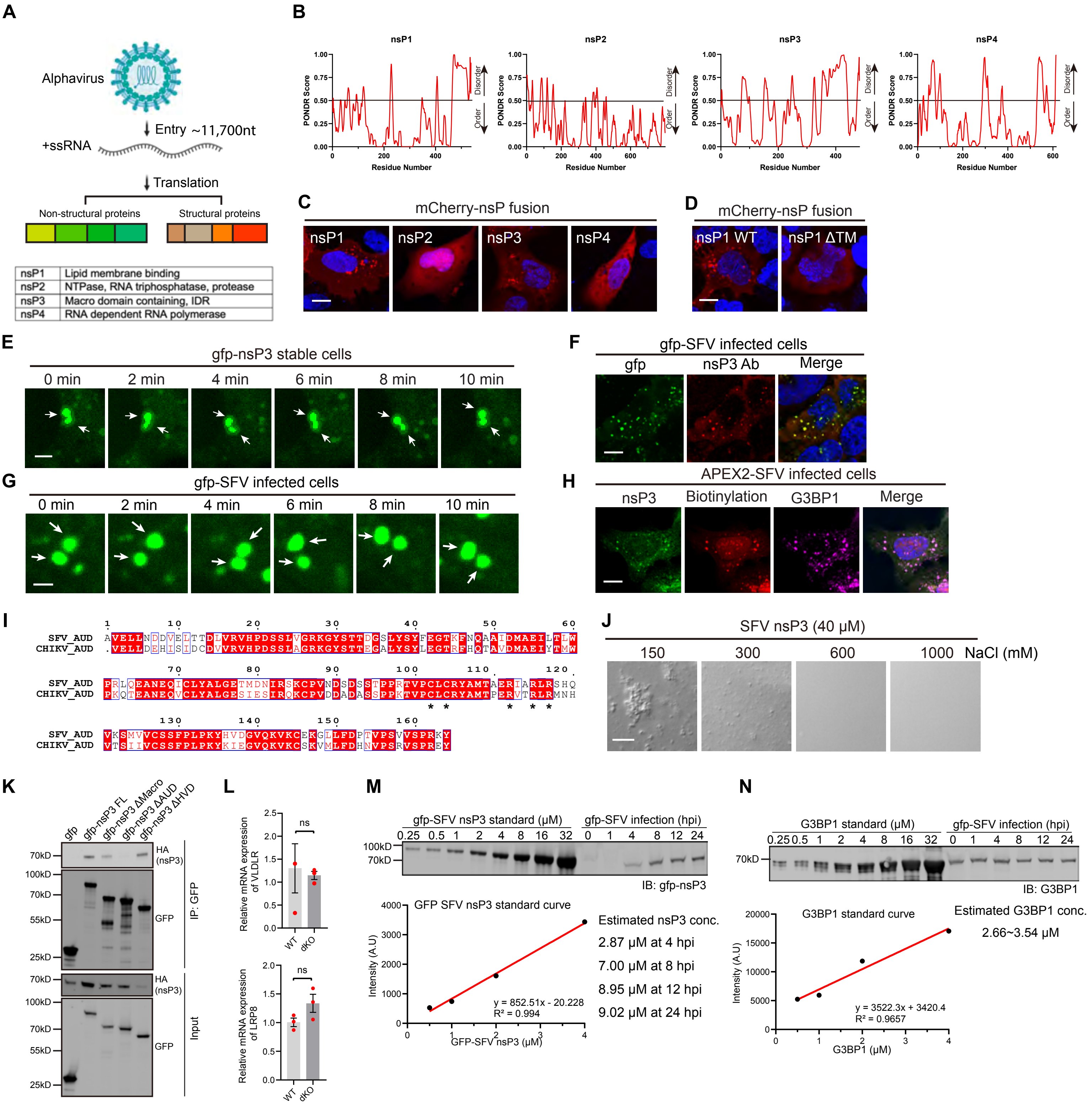

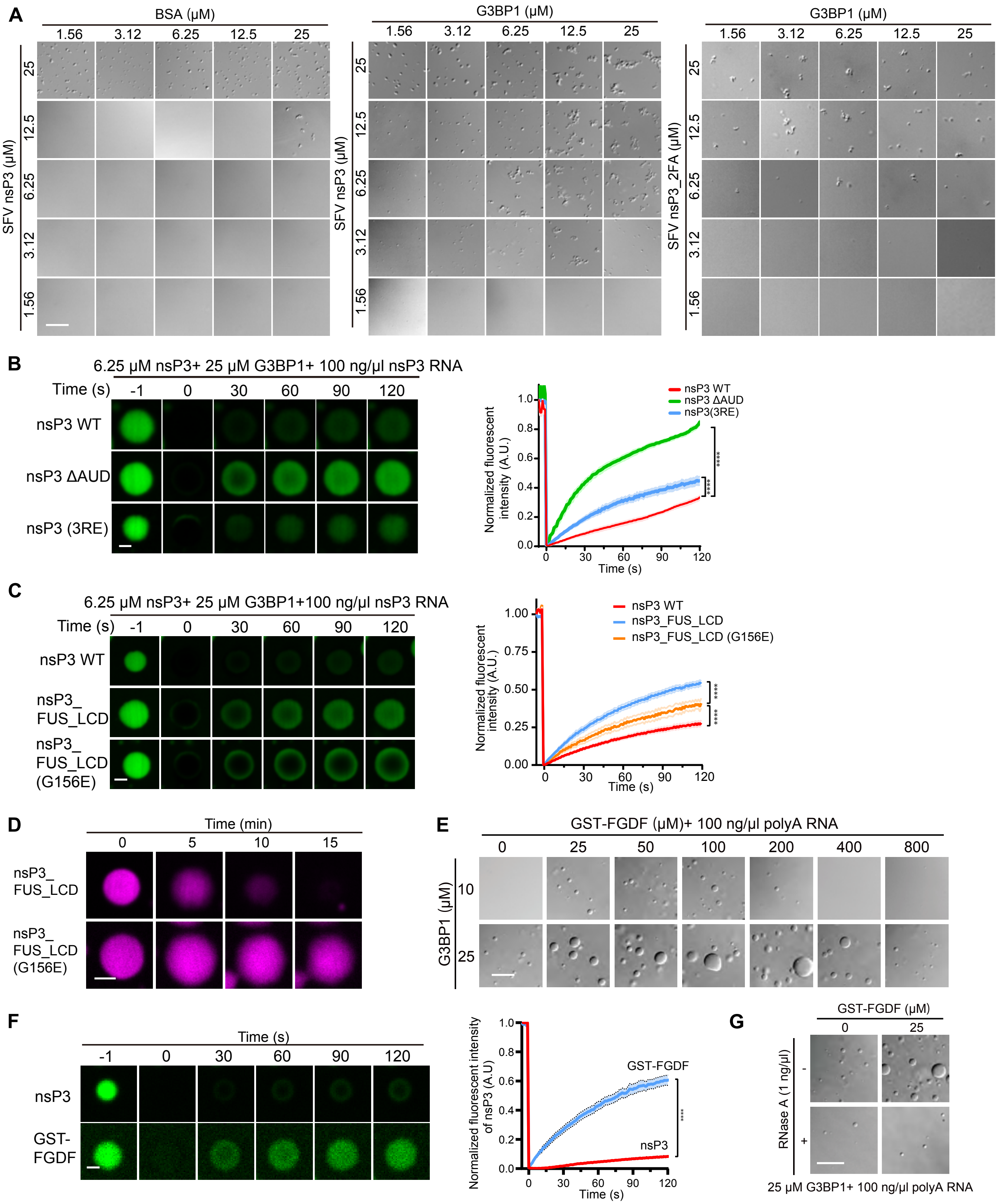

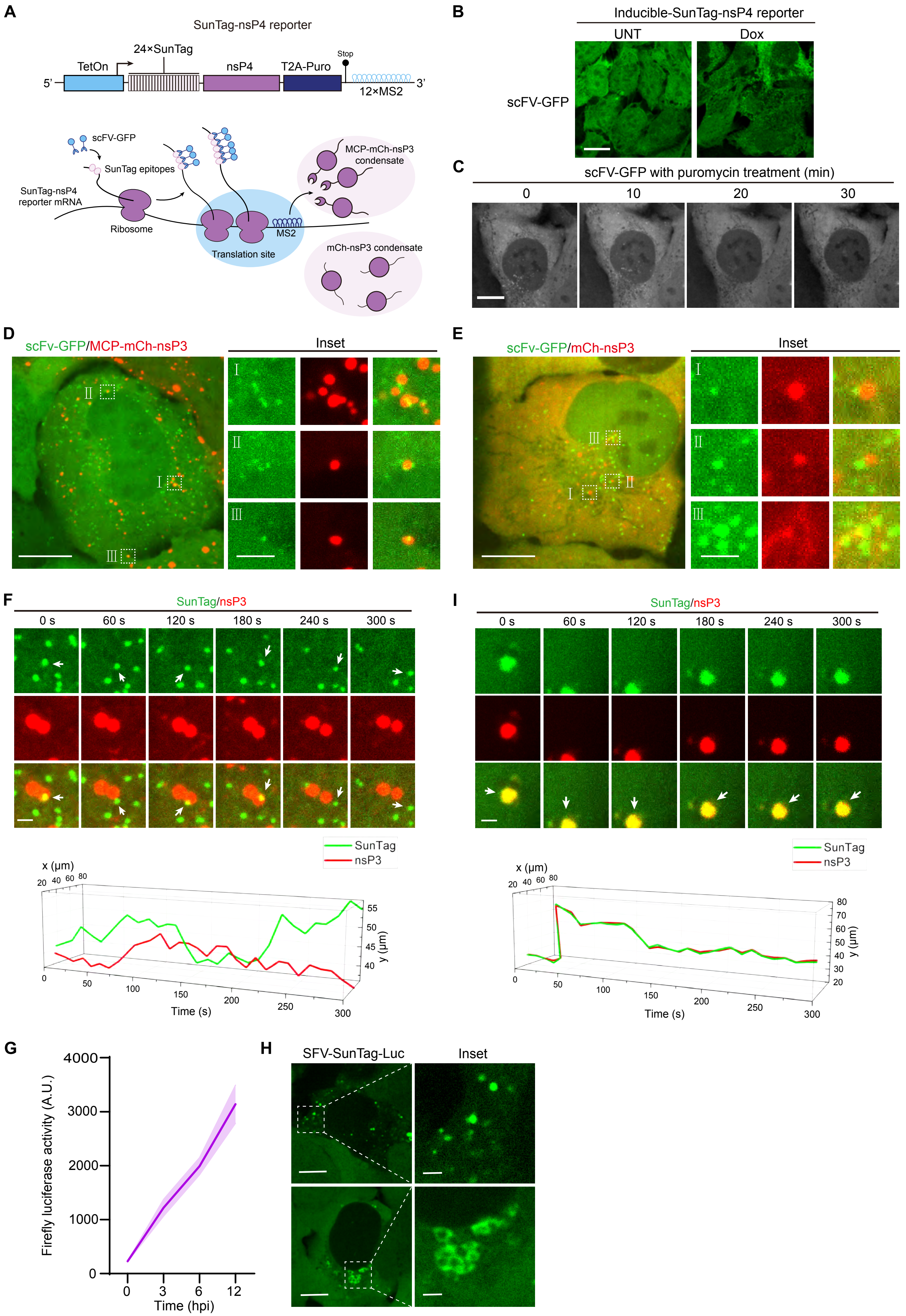

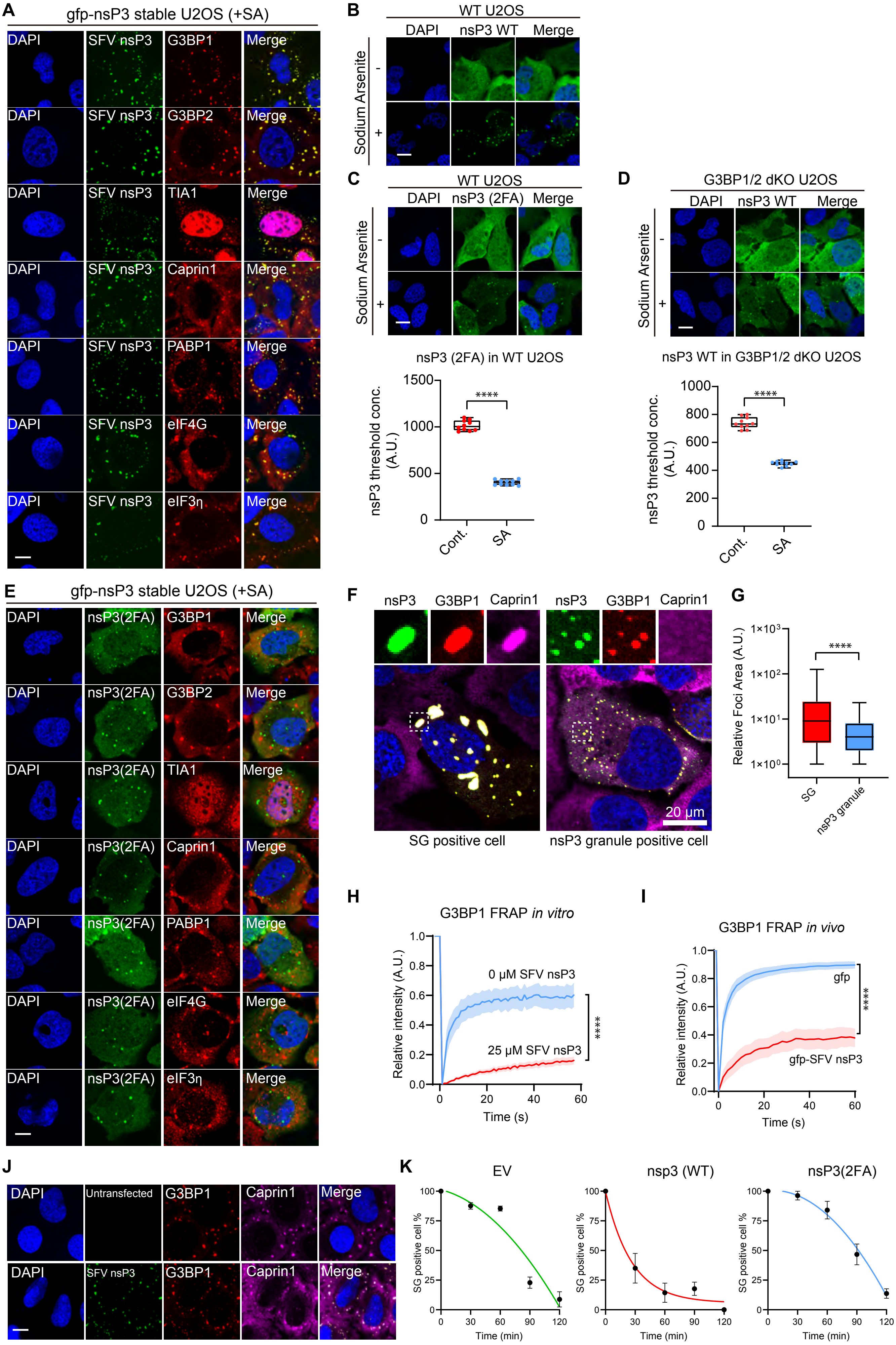

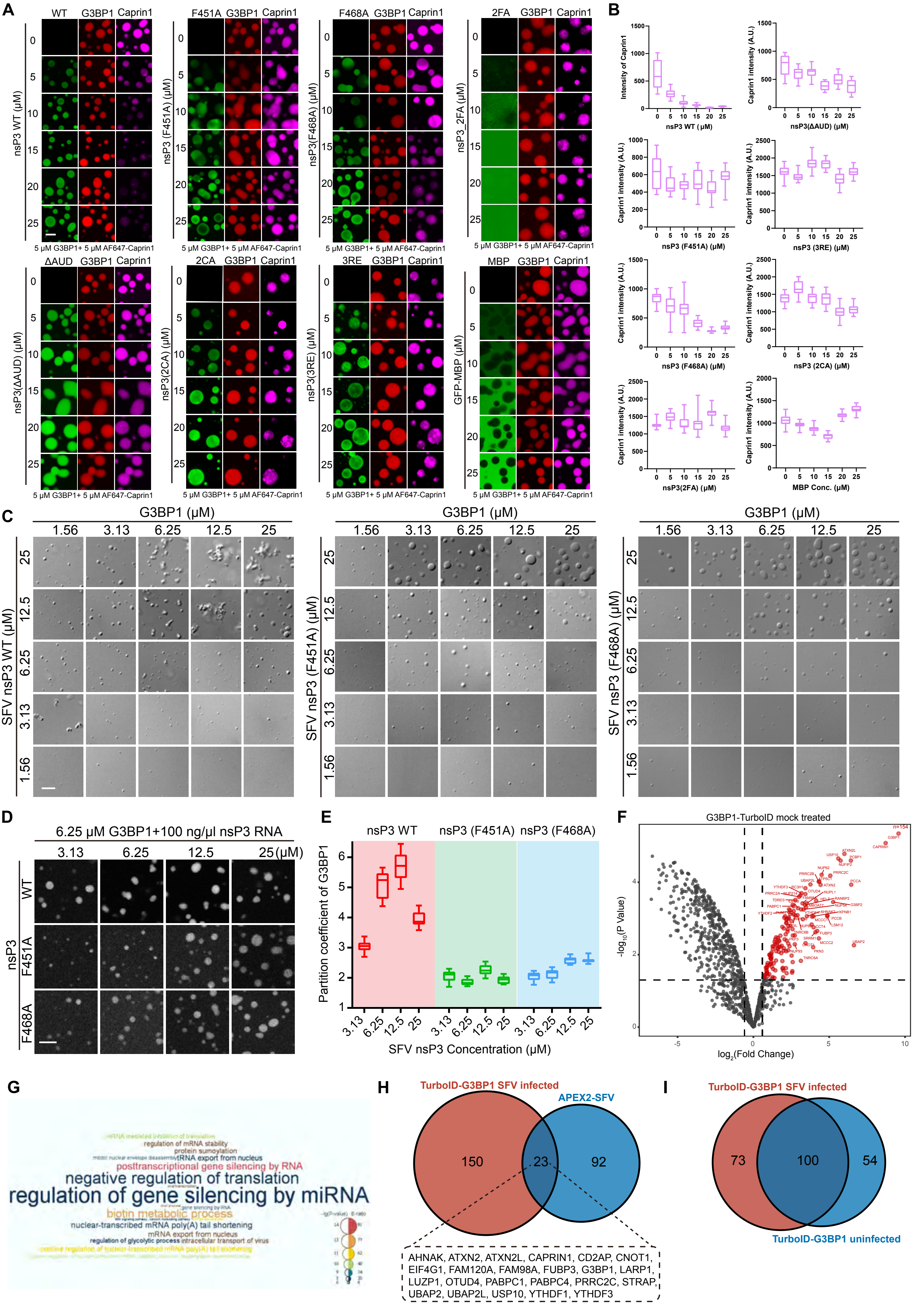

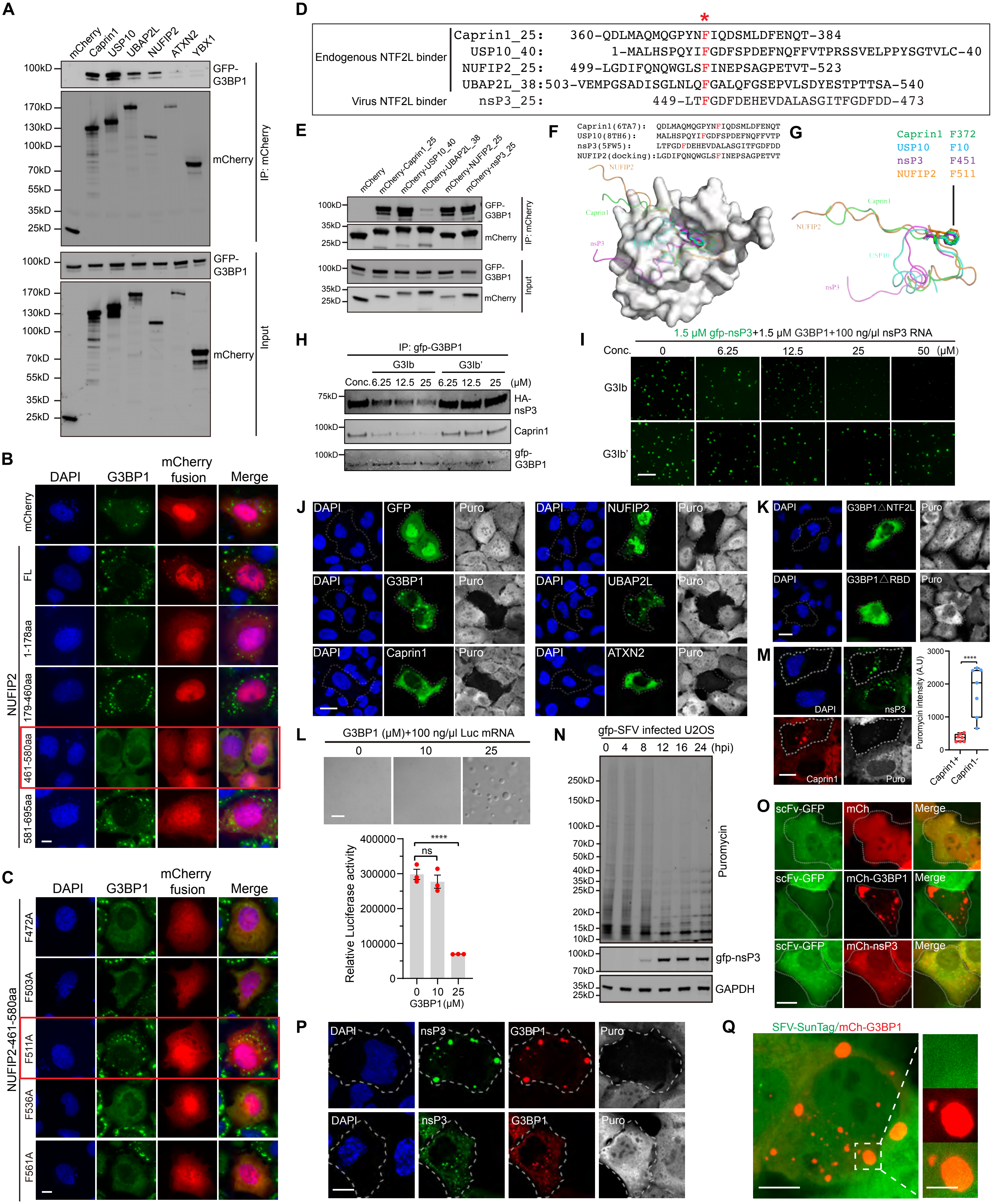

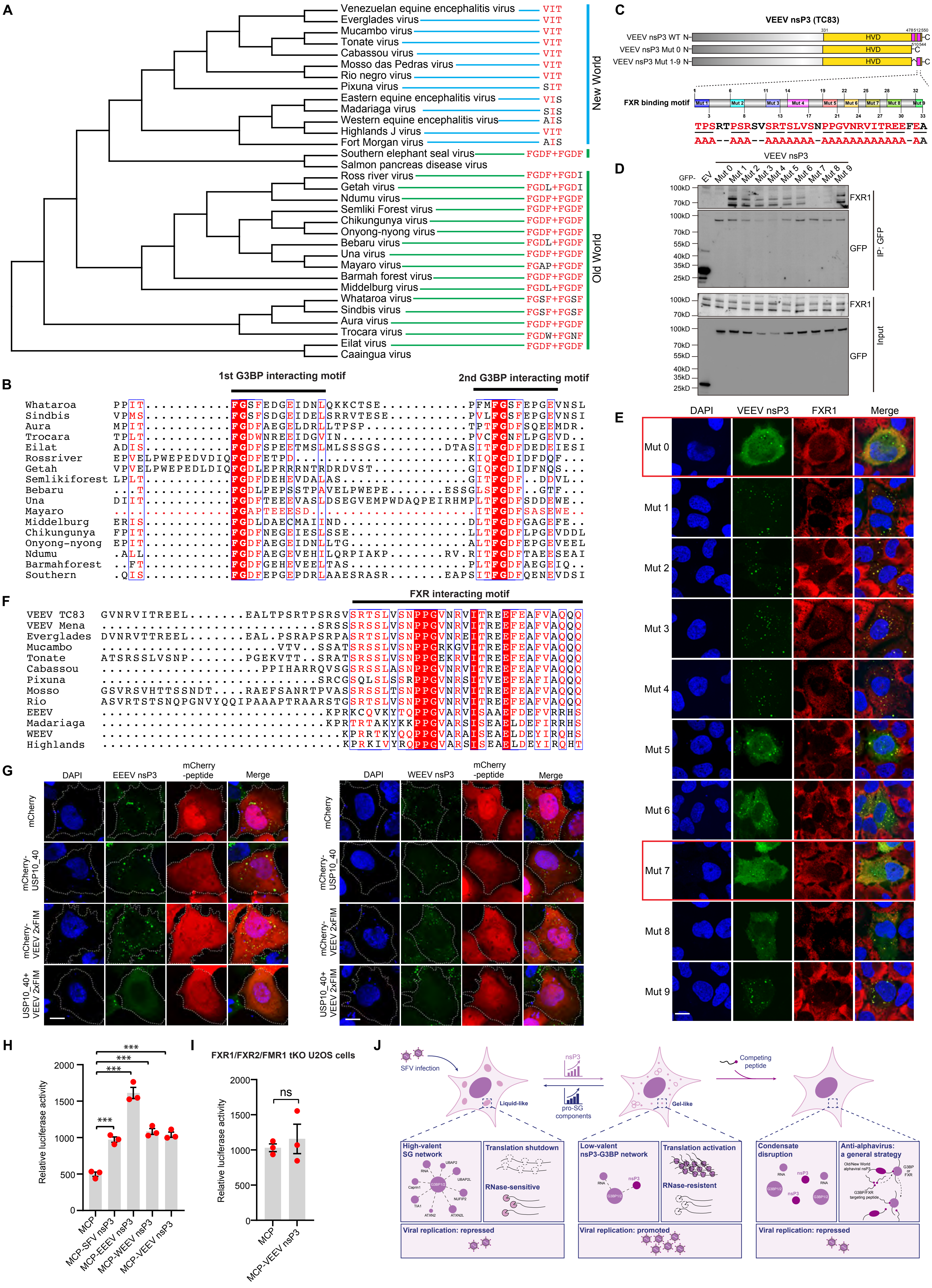
